# Measurement of Accumulation of Antibiotics to *Staphylococcus aureus* in Phagosomes of Live Macrophages

**DOI:** 10.1101/2023.02.13.528196

**Authors:** Joey J. Kelly, Brianna E. Dalesandro, Zichen Liu, Mahendra D. Chordia, George M. Ongwae, Marcos M. Pires

## Abstract

*Staphylococcus aureus* (*S. aureus*) has evolved the ability to persist after uptake into host immune cells. This intracellular niche enables *S. aureus* to potentially escape host immune responses and survive the lethal actions of antibiotics. While the elevated tolerance of *S. aureus* to small-molecule antibiotics is likely to be multifactorial, we pose that there may be contributions related to permeation of antibiotics into phagocytic vacuoles, which would require translocation across two mammalian bilayers. To empirically test this, we adapted our recently developed permeability assay to determine the accumulation of FDA-approved antibiotics into phagocytic vacuoles of live macrophages. Bioorthogonal reactive handles were metabolically anchored within the surface of *S. aureus,* and complementary tags were chemically added to antibiotics. Following phagocytosis of tagged *S. aureus* cells, we were able to specifically analyze the arrival of antibiotics within the phagosomes of infected macrophages. Our findings enabled the determination of permeability differences between extra- and intracellular *S. aureus*, thus providing a roadmap to dissect the contribution of antibiotic permeability to intracellular pathogens.

## Introduction

*Staphylococcus aureus* (*S. aureus*) is a significant human pathogen that can cause a wide range of infections from mild skin infections to severe disease such as sepsis and pneumonia.^1^ The increasing incidence of drug-resistant *S. aureus* infections, particularly methicillin-resistant *S. aureus* (MRSA), is a major concern.^2^ MRSA is resistant to many commonly used antibiotics; this resistance poses a significant challenge in effectively treating MRSA infections and can lead to more severe illness, prolonged hospital stays, and increased healthcare costs. Overall, the increasing incidence of drug-resistant *S. aureus* infections highlights the importance of understanding the factors that impact the efficacy of therapeutics against bacterial pathogens.

In addition to genetic alterations that result in drug resistant phenotypes, *S. aureus* has evolved another mode of persistence in a host organism. Host immune cells, such as macrophages and neutrophils, phagocytose bacteria as an anti-bacterial response.^3, 4, 5, 6^ In turn, immune cells have a variety of potent mechanisms to kill phagocytosed bacteria including the release of reactive oxygen and nitrogen species (ROS and RNS), the production of antimicrobial peptides and enzymes, and the acidification of the phagosome.^7^ Yet, some pathogens continue to remain viable despite these challenging environment.^3, 8^ *S. aureus* that survive inside immune cells can serve as potential reservoirs that can seed reoccurring infections and can persist for days,^4, 9^ a feature that has been demonstrated both *in vitro* and *in vivo*.^10-12^

Mechanistically, antibiotic tolerance by intracellular *S. aureus* has been previously ascribed to a phenotypic switch to small colony variants.^6, 13^ An alternative or complementary mechanism that can promote the persistence of *S. aureus* within immune cells is reduced accumulation of antibiotics to the subcellular organelle housing these cells (**Figure 1A**). Antibiotics must traverse two membrane bilayers (immune cell plasma membrane and phagosome bilayer) to reach the target pathogen relative to extracellular *S. aureus* cells. These additional barriers could potentially diminish the effective concentration of the therapeutic agent within the phagosome, leading to less effective antibacterial activity. Herein, we developed an assay to directly measure the permeability of the antibiotic to the surface of *S. aureus* present in the phagosomes of macrophages (**Figure 1B**). We demonstrated that structural modifications of antibiotics can dictate the abundance of agents within phagocytic vacuoles. Uniquely, these measurements are reflective of antibiotic accumulation at the target bacteria cells and not a total measurement of antibiotic levels in the immune cell.

**Figure 1.**
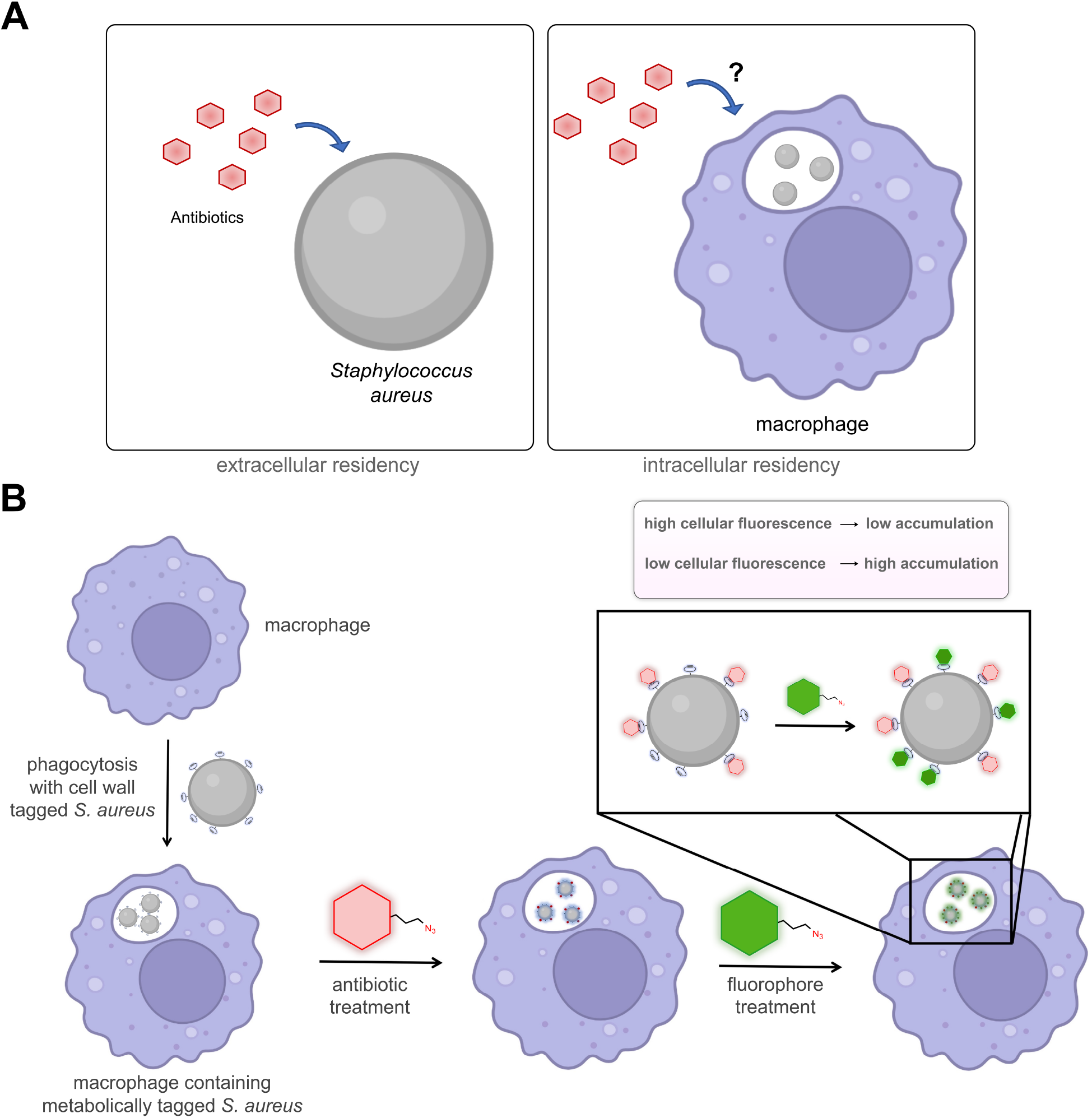
A) Cartoon representation of antibiotic permeability to extracellular *S. aureus* (left) and intracellular *S. aureus* (right). B) Workflow of the assay for measuring accumulation of antibiotics using a competition towards tags within the cell wall of *S. aureus*.

## Results and Discussion

Prior efforts to assess levels of antibiotics in host mammalian cells have most commonly been approximated by their antibacterial activity (minimum inhibitor concentration, MIC).^10, 14^ The main challenges in using MIC determinations as a proxy for the concentration of the antibiotics in the phagosome are that tolerant phenotypes (induced following phagocytosis) can alter MIC values regardless of drug concentration and/or phagosome pH can alter drug efficacy.^15^ Aside from MIC determination, antibiotic concentration measurements have also been made based on the inherent fluorescence of those agents, such as in the case of quinolones^10, 16^ or by introducing a radioactive isotope.^17^ These methods pose clear challenges and limitations including the overall throughput and confounding antibacterial/permeation effects. Moreover, these analyses revealed the antibiotic concentration of the entire immune cell rather than the phagocytic vacuole. The concentration of the antibiotic inside the phagosome could potentially differ from that in the general cytosol; thus, the whole cell concentration may not properly describe the effective level that reaches the target bacterial cells.

Appreciating that intracellular residence can lower the efficacy of antibiotics, a group at Genentech developed an antibody-antibiotic conjugate that eliminated intracellular *S. aureus* in an *in vivo* model of infection. More specifically, a rifamycin derivative was conjugated to an antibody directed at the wall-teichoic acids (WTA) of *S. aureus,* whereby rifamycin was released upon cleavage by intracellular proteases.^11^ Another clever strategy involved the use of a proline-based cell penetrating peptide that was conjugated to kanamycin that targeted intracellular bacteria.^18^ Similarly, vancomycin was conjugated to an arginine-based cell penetrating peptide, which led to its potentiation.^19^ Antimicrobial peptides have also been designed to directly target intracellular bacterial pathogens.^20^ Together, these efforts highlight diverse and promising approaches to targeting intracellular pathogens.^21^ We set out to isolate the impact of intracellular residence to the accessibility of antibiotics by systematically determining the permeability of a panel of antibiotics to the cell wall of intracellular *S. aureus*.

To measure the accumulation of antibiotics, we combined an assay our laboratory recently developed aimed at systematically assessing how flexibility and size of large biopolymers impact their ability to reach the peptidoglycan (PG) surface of *S. aureus* cells^22, 23^ with one that we developed to measure the permeability of molecules into mycobacteria.^24^ In this new iteration (**Figure 1B**), a strained alkyne dibenzocyclooctyne (DBCO) is metabolically installed within the PG of *S. aureus*. DBCO-tagged bacterial cells are then incubated with macrophages to promote their uptake. Treatment of cells with azide-modified antibiotics (azAbx) post infection was expected to result in the reaction between the DBCO and azAbx *via* strain promoted alkyne-azide cycloaddition (SPAAC).^25^ A chase step with an azide-modified fluorophore imprints a signal that is reflective of unmodified DBCO epitopes. Cellular fluorescence measurement should then enable a fluorescent, live-cell analysis of antibiotic accumulation, whereby cellular fluorescence inversely correlates with antibiotic accumulation levels.

The site-specific metabolic installation of a bioorthogonal reactive handle within the PG of *S. aureus* is central to the overall assay. For Gram-positive bacteria, the cell wall is primarily comprised of a thick layer of peptidoglycan (PG) that displays various surface proteins and extracellular matrices; these biomacromolecules are essential to maintain the overall integrity of the cell.^26^ PG is composed of repeating disaccharide units, *N-* acetylglucosamine (GlcNAc) and *N-*acetylmuramic acid (MurNAc), covalently linked to a short stem peptide (L-Ala-D-Glu-L-Lys-D-Ala-D-Ala). The stem peptide strands are crosslinked together by transpeptidase (TP) enzymes to create a dense, mesh-like scaffold that surrounds the cell as a single molecule.^27^ Given that the PG is a critical component to the fitness of the pathogen, many antibiotics target PG biosynthesis enzymes.^28^ Our lab^22, 29, 30^ and others^31^ have extensively demonstrated the ability to metabolically remodel bacterial PG with synthetic analogs of PG precursors. For most of these modifications, the viability and overall cellular structure of the PG remain unaffected. Further, we demonstrated that such analogs can be incorporated *in vivo* into the cell wall of *S. aureus* infected *C. elegans*^30^ and various diverse bacterial species residing in the gut microbiome of a mouse model.^32^

To start, we prepared DBCO-tagged PGs by synthesizing the unnatural amino acid, D-Dap, and conjugating its sidechain with DBCO to yield D-DapD. Incubation of this amino acid in the growth media of *S. aureus* results in the exchange of the terminal D-alanine on the PG stem peptide with the supplemented unnatural amino acid (**Figure S1**). *S. aureus* cells were incubated with culture media containing D-DapD (or its stereocontrol, L-DapD) overnight to allow for incorporation throughout the PG matrix. The modified L-amino acid is not expected to be metabolically processed by the cell wall transpeptidases, and, therefore, no tagging should occur. Bacterial cells were washed to remove unincorporated metabolic tags, followed by incubation with azide-modified rhodamine 110 (R110az), and cellular fluorescence measurement by flow cytometry (**Figure 2A**). Satisfyingly, the fluorescence associated with *S. aureus* cells metabolically labeled with D-DapD were significantly higher than DMSO treated cells (**Figure 2B**). Cells treated with the stereocontrol, L-DapD, exhibited similar fluorescence levels to DMSO treated cells indicating that the DapD was incorporated in the PG scaffold of *S. aureus* in a stereoselective manner. Additionally, incubation with D-DapD did not perturb cell growth (**Figure 2C**). Confocal microscopy showed that the fluorescence pattern was consistent with PG incorporation as the fluorescence remained peripheral and septal (**Figure 2D**).

**Figure 2.**
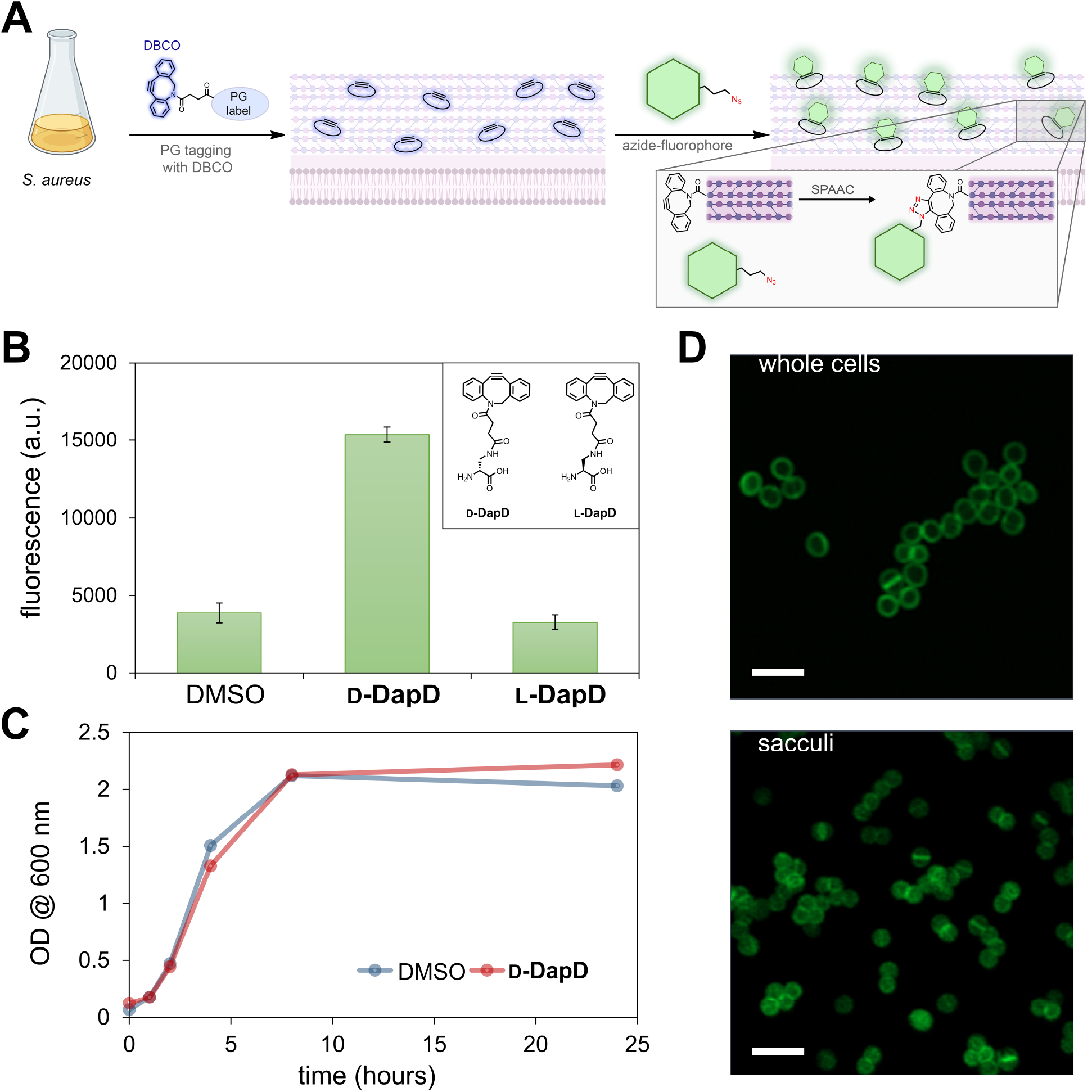
A) Workflow schematic of the PG labeling with a DBCO-displaying agent followed by treatment with an azide-tagged fluorophore. B) *S. aureus* was incubated overnight with 500 μM of D-DapD or L-DapD in tryptic soy broth (TSB) at 37°C. The next day cells were washed 3x with PBS and incubated for an additional 30 min with 25 μM of R110az in PBS at 37°C. Cells were then fixed with 2% formaldehyde in PBS and 10,000 events per sample were analyzed via flow cytometry. C) Measurement of OD_600_ over time for *S. aureus* cells treated with either DMSO or 500 μM of D-DapD in TSB. D) Confocal microscopy analysis of *S. aureus* cells (top) and isolated sacculi (bottom) after treatment with 500 μM of D-DapD in tryptic soy broth. Cells were washed 3x with PBS and incubated for an additional 30 min at 37°C with 25 μM of R110az in PBS before fixation with 2% formaldehyde in PBS. Scale bar = 5 μm. Data are represented as mean ± SD (n= 3) of biological replicates.

We further confirmed DBCO installation on the PG of *S. aureus* by isolating the entire PG scaffold (sacculi) from cells that had been treated with D-DapD followed by R110az. Sacculi isolation was performed using standard methods that disrupt the whole cell, leaving discrete and intact units of sacculi that are similar in size.^33^ Isolation of sacculi with the DBCO tag was not performed due to the chemical instability of the triple bond of DBCO in the isolation conditions (acid, high temperatures). Instead, we isolated the stable triazole product formed after the SPAAC reaction. Flow cytometry analysis of the fluorescently tagged sacculi (SaccuFlow^33^) was performed wherein the sacculi was found to be highly fluorescent (**Figure S2**). Sacculi isolated from DMSO-treated *S. aureus* cells displayed minimal fluorescence levels. As expected, confocal microscopy analysis revealed that the fluorescently labeled sacculi were similar in size and shape compared to whole cell *S. aureus* while also retaining similar labeling patterns (**Figure 2D**). Together, these results established that tagging of the sacculi was successful and the DBCO sites could be specifically labeled by treatment with an azide-modified fluorophore.

We next set out to test labeling conditions to optimize the dynamic range of the assay. Following the same workflow as described before, *S. aureus* cells were grown overnight in the presence of increasing concentrations of D*-*DapD, washed, treated with R110az, and analyzed by flow cytometry. As expected, the cellular fluorescence levels of *S. aureus* grown with D*-*DapD increased in a concentration-dependent manner (**Figure S3A**); this indicates that more D*-*DapD was covalently linked into the growing PG scaffold as more amino acid was supplemented into the growth media. From these results, the D*-*DapD concentration of 500 μM was selected for subsequent experiments. We next evaluated the impact of fluorophore concentration on the cellular fluorescence levels. *S. aureus* cells were labeled D*-*DapD followed by incubation with increasing concentrations of R110az, which resulted in increasing cellular fluorescence levels, as expected (**Figure S3B**). The selected parameters of D*-*DapD and R110az concentrations were based on the large dynamic range that these conditions afforded while reducing reagent consumption and the exposure to non-native molecules. To ensure the timeframe of the assay on the surface of *S. aureus* will enable completion of the SPAAC reaction, a time course of R110az labeling was performed, which indicated that the R110az reaction mostly plateaued within 30 minutes (**Figure S3C**). Lastly, to investigate the permeation effects of the fluorophore itself, we examined two additional azide fluorophores: fluorescein azide (Flaz) and coumarin azide (Coaz). These dyes have different physiochemical properties (charges/molecular weight) than R110az and could potentially result in superior signal-to-noise ratios. Although inferior to R110az, both were demonstrated to be compatible (**Figure S3D**) as indicated by significant fluorescence signals over cells in the absence of the metabolic label.

We then evaluated the ability of the assay to report on the accessibility of FDA-approved antibiotics modified with azide tags to the PG of *S. aureus* in phagocytic vacuoles. We synthesized an azAbx panel to make them compatible with our assay workflow (**Figure 3A**). Azides were installed on parts of the antibiotic that were found to minimally perturb their structure (Table 1). The azAbx included were rifampicin, erythromycin, linezolid, and ciprofloxacin, which are treatment options for *S. aureus* infections.^34^ The selected antibiotics have chemical compositions that range in hydrophobicity, polarity, size, and charge to examine how these factors affect their ultimate accumulation into phagosomes. Furthermore, we also tested two variations of ciprofloxacin: one containing the azide modification on the piperazine group that has an overall net negative charge and another that has an additional ester modification to return the molecule to a net neutral charge. This modification was done to evaluate the impact of net charge on the permeability of antibiotics to the phagosome. In the pulse step, DBCO-tagged cells were treated with the azAbx. Antibiotics that reach the PG are expected to form a stable and irreversible bond with DBCO. In the chase step, R110az reacts with the remaining intact DBCO sites on the PG. Therefore, antibiotics that readily reach the bacterial pathogens should result in low cellular fluorescence levels (**Figure 1B**).

**Figure 3.**
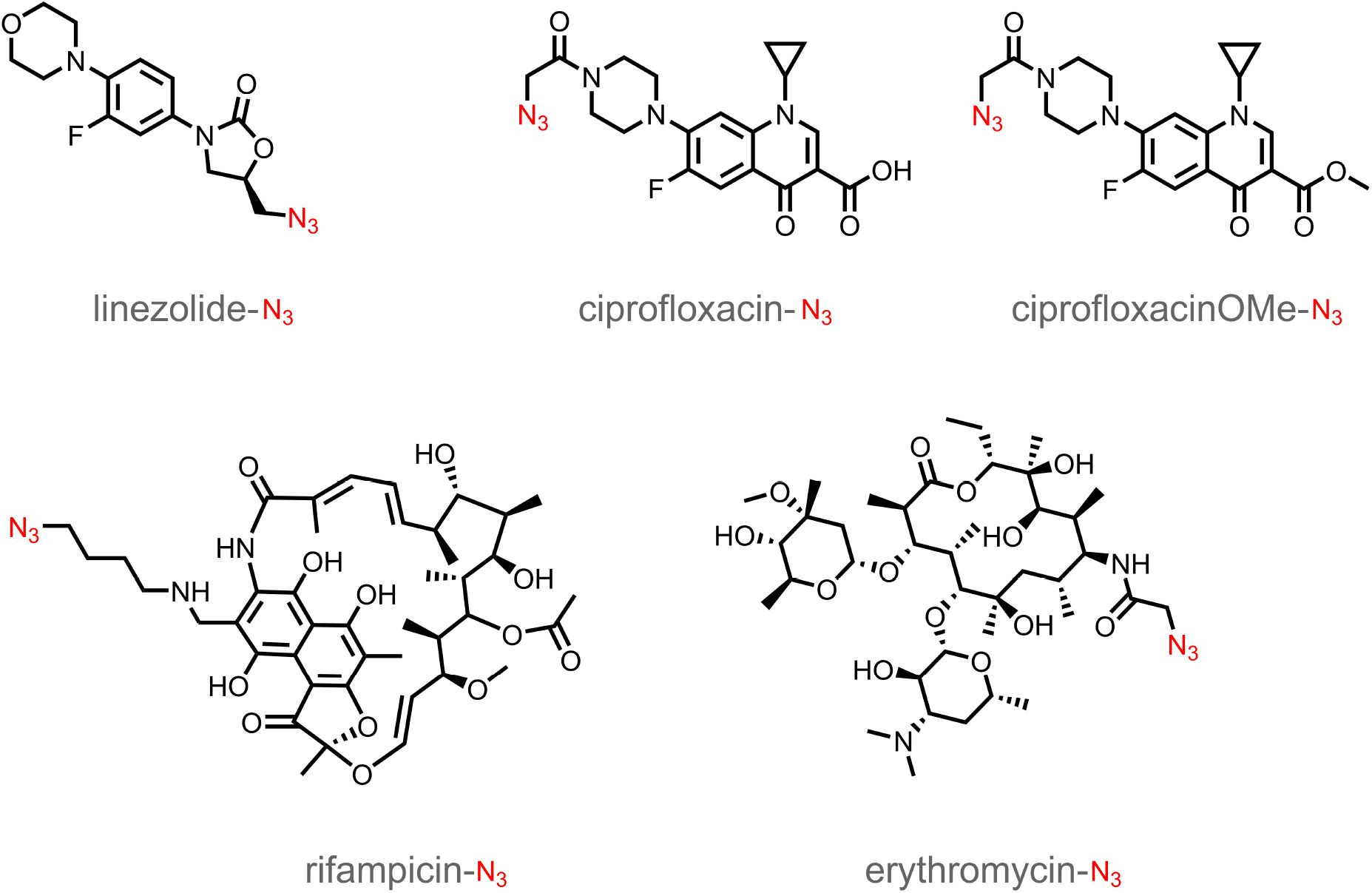
Chemical structures of azide modified antibiotics.

To establish the baseline reactivity level of the azAbx without a membrane-permeability barrier, *S. aureus* cells were metabolically tagged with D-DapD and incubated with each individual member of the azAbx library. All azAbx were tested at 25 μM based on our findings using R110az. Following incubation with the azAbx, cells were washed, treated with R110az, and cellular fluorescence was analyzed by flow cytometry (Figure 4A). From the data observed, it was found that nonpolar antibiotics, such as linezolid and rifampicin, were able to sieve through the PG and fully react with the DBCO modified PG within 2 h, as indicated by a low fluorescence signal. The small size of linezolid and the flexible backbone of rifampicin could be, at least in part, the reason for these observations.

Conversely, the increased rigidity and polarity of ciprofloxacin and erythromycin could play a role in hindering the permeation of these molecules through the PG; to this end, full reaction was not observed until nearly 8 h (**Figure 4A**). When extracellular *S. aureus* was treated with untagged antibiotics, we observed no change in cellular fluorescence levels (**Figure S5**). These results suggest that the change in fluorescence observed in Figure 4A are not related to bacterial cell lysis/death. Interestingly, a similar pattern of permeation was observed for the two ciprofloxacin derivatives, indicating that masking the net negative charge of ciprofloxacin-Nhad negligible effects in terms of reaching DBCO sites within PG of *S. aureus*.

**Figure 4.**
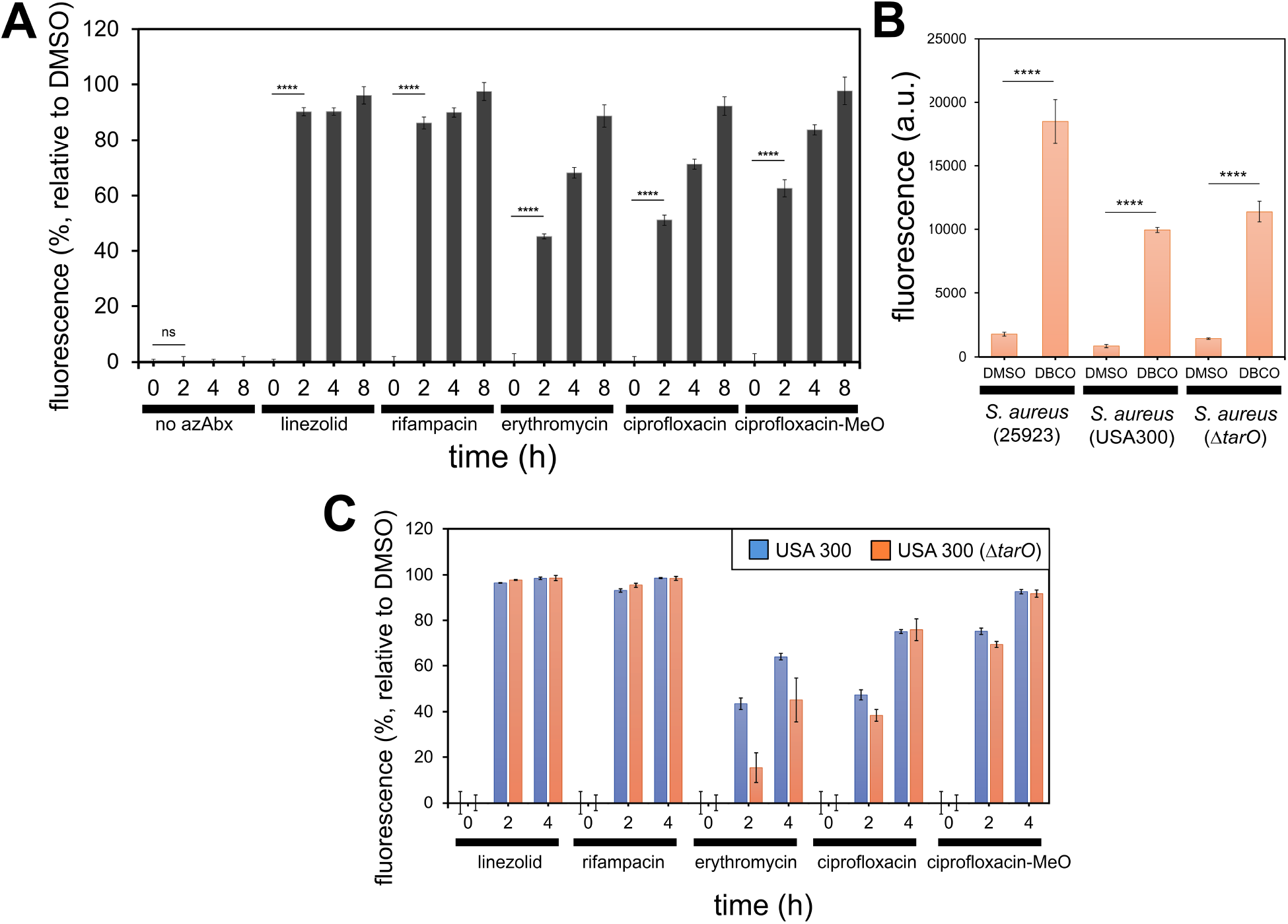
A) *S. aureus* was incubated overnight with 500 μM of D-DapD in TSB at 37°C. The next day cells were washed 3x with PBS and incubated with 25μM azABx in PBS for indicated time points at 37°C. Following azABx incubation, the cells were resuspended in 25 μM of R110az in PBS for 30 min at 37°C and fixed with 2% formaldehyde in PBS before 10,000 events per sample were analyzed via flow cytometry. B) *S. aureus* 25923, USA 300, and a WTA deletion strain, *tarO*, were incubated overnight with 500 μM of D-DapD in TSB at 37°C. The following day the cells were washed 3x with PBS and incubated with 25 μM R110az in PBS for 30 min at 37°C. Cells were then fixed with 2% formaldehyde in PBS and 10,000 events per sample were analyzed via flow cytometry. (C) *S. aureus* cells (USA300 and *tarO*) were incubated with 500 μM of D-DapD overnight, washed 3x with PBS, and followed by incubation with 25 μM of azABx in PBS for indicated time points. Following azAbx incubation, the cells were resuspended in 25 μM of R110az in PBS for 30 min, fixed in 2% formaldehyde in PBS, and 10,000 events per sample were analyzed via flow cytometry. Data are represented as mean ± SD (n= 3) of biological replicates. *P*-values were determined by a two-tailed *t*-test (**** *p* < 0.0001, ns = not significant).

To further ensure whether that the SPAAC reaction between the azAbx and D-DapD was complete within the timeframe of the assay, we modified amine-terminated beads to display DBCO epitopes (**Figure S5A**). As these beads are compatible with flow cytometry, this approach provides a parallel workflow to the live cell analysis but without the potential barrier of the matrix-like PG scaffold. Our results showed that the antibiotics in this panel readily react in the absence of a complex cell wall structure, as indicated by background fluorescence levels observed for all five azAbx within 2 h (**Figure S5B**). These results clearly suggest that the PG itself could pose as a permeation barrier. Sieving through the crosslinked PG has been ascribed to play a role in the activity of antimicrobial peptides^35^ and components of the human immune system (e.g., lysozyme and antibodies).^36, 37^ Early work showed that molecular sieving was operative within isolated sacculi with synthetic polymers^38^ but systematic studies had not been carried out in live bacterial cells using antibiotics. Sieving within the PG scaffold is likely to be dependent on multiple factors, including the level of crosslinking, which can vary across strains and species.^39^ Given that the estimated crosslinking levels in *S. aureus* can range from 80 to 90%, it is probable that sieving could be particularly prominent.^40^

To demonstrate the adoptability of the assay across other strains of *S. aureus*, we also performed a similar analysis with MRSA. Satisfyingly, treatment of USA300 *S. aureus* with D-DapD resulted in a high fluorescence signal when incubated with R110az, indicating that D-DapD was incorporated into the PG layer of a methicillin-resistant strain and readily reacted with an azide molecule (**Figure 4B**). Further, we went on to investigate azAbx accessibility to USA300 PG. After incubating D-DapD labeled USA300 with the azAbx, a similar pattern was observed for all azAbx compared to methicillin sensitive *S. aureus* (25923, MSSA), as evidenced by a significant decrease in fluorescence signal compared to DMSO treated cells (**Figure 4C**) after incubation with the azAbx for 4 h. This indicates that the azAbx were able to sieve through the USA300 PG layer.

Further, we aimed to investigate the contribution that PG-anchored, brush-like polysaccharides play on permeability. The surface of *S. aureus* is coated with wall teichoic acids (WTAs), that are anionic glycopolymers known to sterically hinder biomolecules from reaching the surface of the PG.^22, 36, 41^ Our group recently described how WTA can be a major determinant of surface accessibility to large polymers (polyprolines and PEG-based), which was determined by also using a combination of PG tags and SPAAC.^22^ Analogously, we tested whether WTAs may alter the accessibly of azAbx, which are considerably smaller, to the PG layer. Our data showed no significant difference between the USA300 parental strain and *tarO* deletion strain, which cannot biosynthesize WTA on the surface of *S. aureus* (**Figure 4B,C**). These results indicate that WTAs do not appear to hinder permeation of smaller molecules to the PG matrix. To the best of our knowledge, this had not been shown before.

With benchmarks established for extracellular *S. aureus*, we next sought to test the reportability of molecule accumulation inside phagocytic vacuoles of macrophages (**Figure 1B**). Given the selectivity of the SPAAC reaction, we reasoned that it was possible to isolate accumulation measurements (decoupled from biological activity) inside this cellular compartment. We initially optimized the role of the reporter dye in the absence of azAbx. *S. aureus* were tagged with D-DapD overnight and then incubated with macrophages to promote their uptake. Macrophages containing DBCO-tagged *S. aureus* (or unmodified *S. aureus*) were treated with either 5 μM or 25 μM of R110az (**Figure 5A**). Minimal differences in cellular fluorescence levels were observed at the lower concentration. Upon increasing the dye concentration, a marked difference emerged between DBCO-tagged cells and untagged cells. Additionally, a lower multiplicity of infection (MOI) resulted in poorer differences in cellular fluorescence (**Figure S5**). It was determined that MOI of 100 could detect significant changes in R110az uptake for D-DapD *S. aureus* over the unlabeled DMSO control, as indicated by a significant change in fluorescence. Confocal microscopy at two different time points revealed that fluorescence levels were elevated in areas that were consistent with the shape of bacterial cells inside macrophages (**Figure 5B**). This indicates that the bacterial cells appear to remain intact and not lysed, as expected. Wheat germ agglutinin (WGA) staining was used to broadly stain glycoproteins on the cell membrane but also stains the PG bacterial cells.

**Figure 5.**
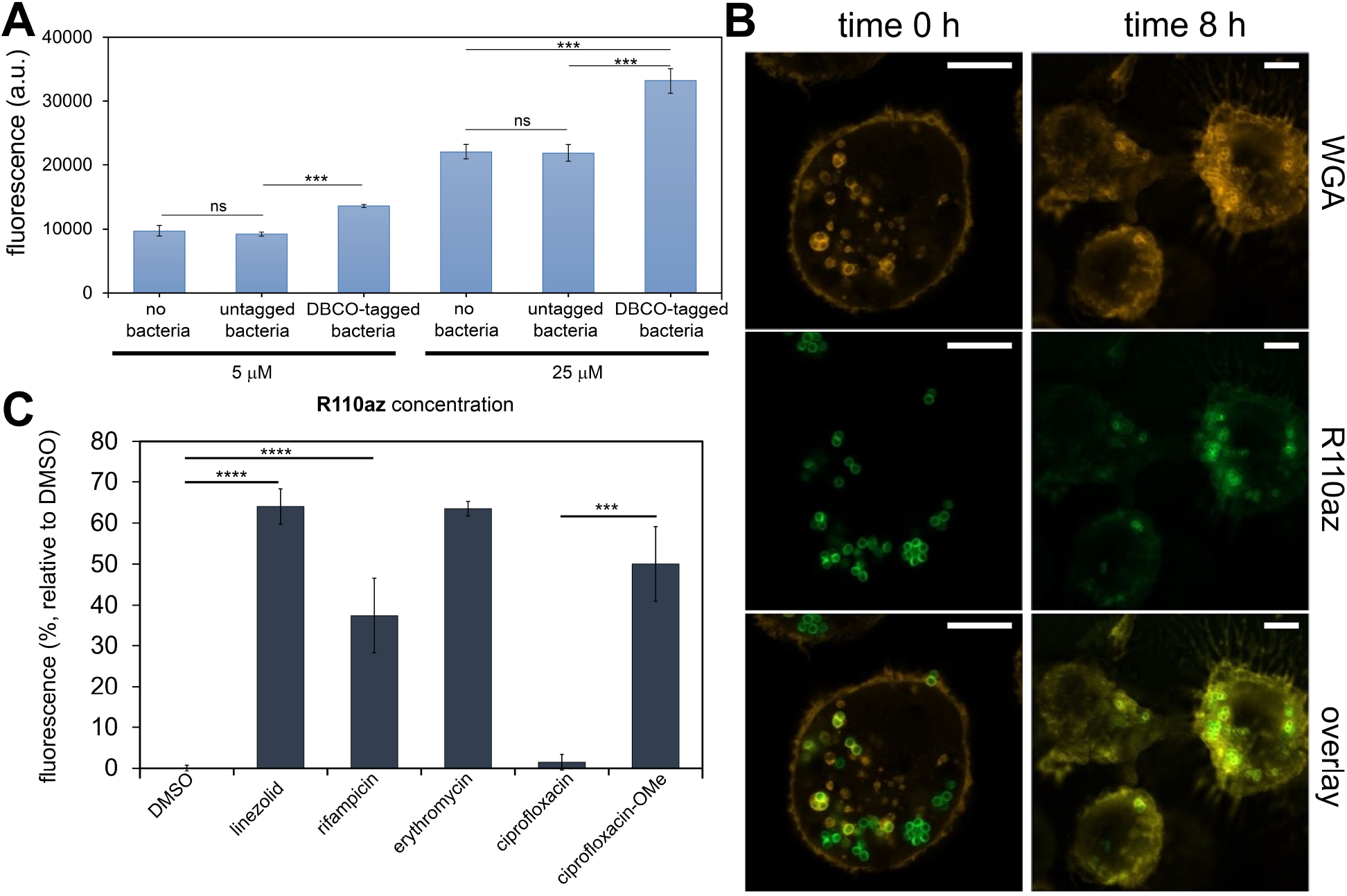
A) *S. aureus* was grown overnight with 500 μM of D-DapD in TSB at 37°C. The following day the *S. aureus* was washed 3x with PBS and infected J774A macrophages in DMEM containing no antibiotics at an MOI of 100 for 1 h in a humidified incubator at 37°C and 5% CO_2_. Following bacterial uptake, macrophages were incubated with varying concentrations of R110az in PBS for 30 min, fixed with 4% formaldehyde in PBS, and 10,000 events per sample were analyzed via flow cytometry. B) Confocal analysis of D-DapD infected J774A macrophages. *S. aureus* was labeled overnight with 500 μM of D-DapD and then immediately reacted with 25 μM of R110az. Fluorescently labeled *S. aureus* was infected into J774A macrophages at MOI 100 in a humidified incubator at 37°C and 5% CO_2_. Immediately following bacterial uptake, time 0 h samples were fixed and analyzed by confocal microscopy. Time 8 h samples were resuspended in PBS for 8 h, fixed with 4% formaldehyde in PBS, and analyzed by confocal microscopy. Macrophage membranes were labeled with 5 μg/mLwheat germ agglutin (WGA) tetramethylrodamine. Scale bar = 10 μm C) *S. aureus* was grown overnight with 500 μM of D-DapD in TSB at 37°C. The following day the *S. aureus* was washed 3x with PBS and infected J774A macrophages in DMEM containing no antibiotics at an MOI of 100 for 1 h in a humidified incubator at 37°C and 5% CO_2_. After that, the azAbx library was incubated with bacteria infected macrophages for 8 h before being incubated with 25 μM of R110az, fixed with 4% formaldehyde in PBS, and 10,000 events per sample were analyzed by flow cytometry. Data are represented as mean ± SD (n= 3) of biological replicates. *P*-values were determined by a two-tailed *t*-test (* *p* < 0.05, *** *p* < 0.001, ns = not significant).

Finally, the accumulation levels of the molecules in the azAbx panel were tested. After macrophage phagocytosis of D-DapD labeled *S. aureus*, cells were incubated with the azAbx panel over an 8 h period (**Figure 5C**). When internalized *S. aureus* labeled with D-DapD was incubated with linezolid and erythromycin, fluorescence levels were similar to those of extracellular *S. aureus*; these results are suggestive that these antibiotics can readily reach phagocytic vacuoles. Conversely, ciprofloxacin appeared unable to access the phagosome, as evidenced by indistinguishable fluorescence levels between ciprofloxacin incubated cells and those in the absence of an azAbx. We reason that the net negative charge of ciprofloxacin-N_3_ could play a role in reducing the permeability of this antibiotic across the membrane bilayers. To test this, a derivative was also made in which the carboxylic acid was masked as a methyl ester. Our data showed that a marked improvement in permeability was observed, as the fluorescence for intracellular *S. aureus* treated with ciprofloxacin-MeO was significantly lower than those treated with ciprofloxacin.

The masking of the carboxylic acid of ciprofloxacin (and other fluoroquinolones) as an ester has been explored in the past.^42^ Moreover, our results may indicate that performing phenotypic susceptibility testing *via* MIC against intracellular pathogens may be biased by the inability of an antibiotic to reach the phagosome. We demonstrated that masking groups known to hinder membrane permeability, i.e. ciprofloxacin-MeO, improved antibiotic accumulation to the phagosome, emphasizing the need to develop strategies to improve the permeability profiles of existing antibiotic classes. Therefore, our results highlight the importance of taking antibiotic accessibility to the phagosome into consideration when evaluating drug effectiveness against intracellular pathogens. Furthermore, we successfully demonstrated the applicability of our assay to empirically analyze the permeability of antibiotics to the surface of intracellular *S. aureus.* We anticipate that this platform can serve as foundational precedence to address the permeability of other classes of antibiotics to aid in developing effective treatments for persisting intracellular *S. aureus* infections.

Additionally, we applied our assay workflow to another intracellular pathogen, *Streptococcus pyogenes* (*S. pyogenes*), to establish that our concept can be translated to other bacterial pathogens. As such, *S. pyogenes* was labeled overnight with D-DapD and incorporation levels were assayed by incubation with R110az. Our results demonstrated a significant increase in cellular fluorescence compared to unlabeled *S. pyogenes*, indicating successful metabolic labeling and SPAAC on the surface of *S. pyogenes* (**Figure S7A**). As expected, azide-tagged linezolid treatment led to a reduction in cellular fluorescence. Next, D-DapD labeled *S. pyogenes* was incubated with macrophages to enable cell uptake. Cells treated with linezolid led to a decrease in cellular fluorescence, indicating that the mammalian cell and phagosome bilayer did not hinder linezolid permeation to intracellular *S. pyogenes* (**Figure S7B**). Overall, our results indicate that our assay workflow can be applied to other intracellular pathogens to aid in the assessment of antibiotics accumulation in phagocytic vacuoles.

## Conclusion

We demonstrated that the assay we developed is a suitable approach to assess the permeability of antibiotics to the surface of extracellular Gram-positive pathogens and in the phagocytic vacuoles of macrophages. A select group of FDA approved antibiotics were successfully modified with an azide handle to covalently react with DBCO moieties embedded within the PG of *S. aureus* to give a direct read out of antibiotic accumulation. Understanding the ability of antibiotics to penetrate both the mammalian cell membrane and phagosome is critical in terms of treating intracellular *S. aureus* infections. With our approach, we were able to identify that the overall efficacy of certain classes of antibiotics against intracellular *S. aureus* may be largely impacted by the membrane barrier provided by the phagocytic cell. Therefore, it is critical to consider additional membrane components when it comes to treating intracellular bacterial infections. We anticipate our assay will be valuable in distinguishing the root cause of antibiotic resistance in intracellular pathogens.

## Acknowledgement

This study was supported by the NIH grant GM124893-01 (M.M.P.).

## Supporting Information

Additional figures, tables, and materials/methods are included in the supporting information file.

## Notes

### Competing Interest Statement

The authors have declared no competing interest.

### Summary of Updates

Additional data was obtained and is now included.

