## Supporting Information for "Measurement of Accumulation of Antibiotics to *Staphylococcus aureus* in Phagosomes of Live Macrophages"

### Table of Contents


| | |
|---|---|
| **Supporting Figures** | **S4-S9** |
| Figure S1: Mode of incorporation of single amino acids into the 5th position of the PG stem peptide | S4 |
| Figure S2: Flow cytometry analysis of whole bacterial cells and sacculi | S5 |
| Figure S3: *in vitro* PAC-MAN optimization | S6 |
| Figure S4: Minimum inhibitory concentration (MIC) values of azAbx | S7 |
| Figure S5: Synthesis of DBCO- modified polystyrene beads and azAbx competition to DBCO- modified polystyrene beads | S8 |
| Figure S6: Multiplicity of infection (MOI) assay | S9 |
| Figure S7: Application of assay workflow to *S. pyogenes* | S10 |
| Table S1: Minimum Inhibitory Concentration (MIC) of azAbx library | S11 |
| **Methods and Materials** | **S12-S15** |
| Materials | S12 |
| Bacterial cell culture | S12 |
| Mammalian cell culture | S12 |
| Labeling of whole cell bacteria with D-DapD | S12 |
| Sacculi isolation of D-DapD labeled *S. aureus* | S12 |
| azAbx competition to D-DapD labeled whole bacterial cells | S13 |
| Non-Azide competition of D-DapD labeled whole bacterial cells | S13 |
| MIC assay of azAbx against *S. aureus* | S13 |
| Confocal microscopy analysis of whole cell bacteria and sacculi | S14 |
| DBCO modification of polystyrene beads | S14 |
| Competition of azAbx to DBCO modified polystyrene beads | S14 |
| azAbx competition to D-DapD labeled intracellular *S. aureus* | S14 |
| Confocal microscopy of D-DapD labeled intracellular *S. aureus* | S15 |

| Synthesis and Characterization | S16-S26 |
|---|---|
| Scheme S1: D-DapD | S16 |
| Scheme S2: Rifampicin-azidobutane | S19 |
| Scheme S3: Erythromycin-azidoacetamide | S21 |
| Scheme S4: N-azidoacetyl-Ciprofloxacin | S23 |
| Scheme S5: Ciprofloxacin-azidoacetyl methyl ester | S25 |

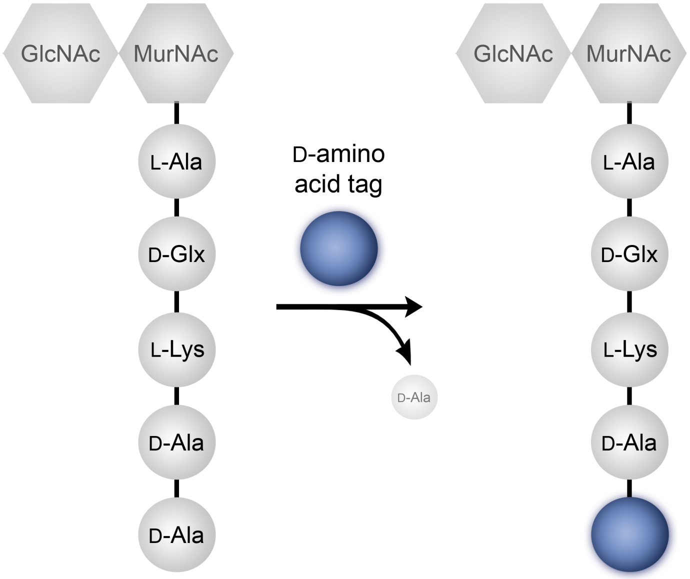


**Figure S1.** Mode of incorporation of single amino acids into the 5th position of the peptidoglycan stem peptide.

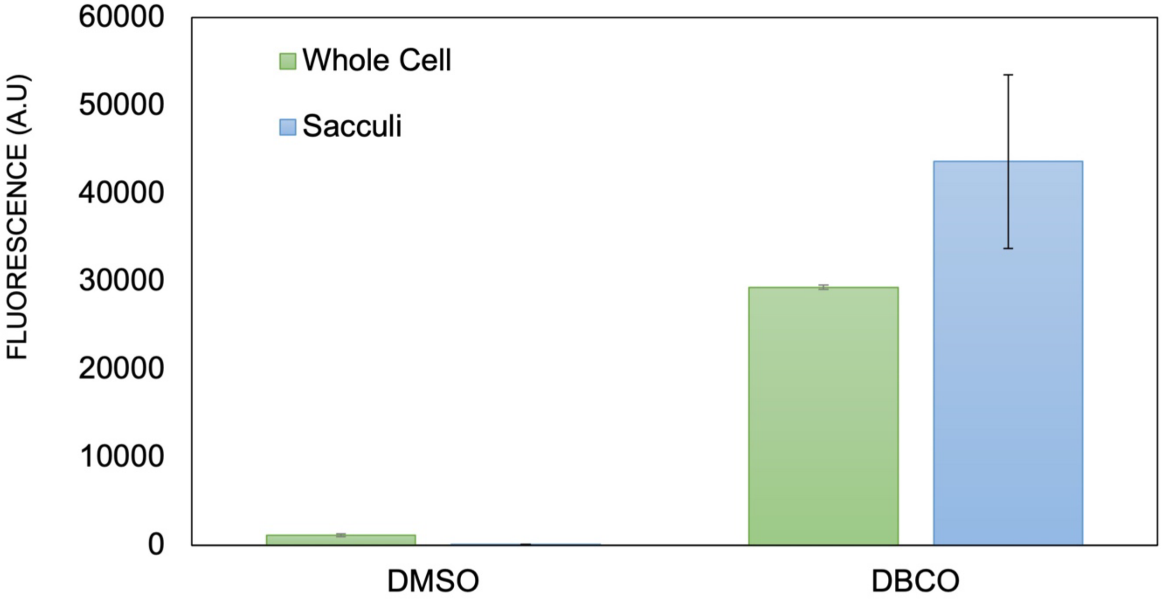


**Figure S2. Flow cytometry analysis of whole bacterial cells and isolated bacterial sacculi.** *S. aureus* 25922 were labeled overnight with 500 μM of D-DapD followed by incubation with 25 μM of R110az. Whole cell samples were analyzed by flow cytometry. Following incubation with R110az, both DMSO-treated and DBCO-treated *S. aureus* sacculi were isolated and analyzed by flow cytometry. 10000 events were recorded for each condition. Data are represented as mean ± SD of biological replicates (n= 3).

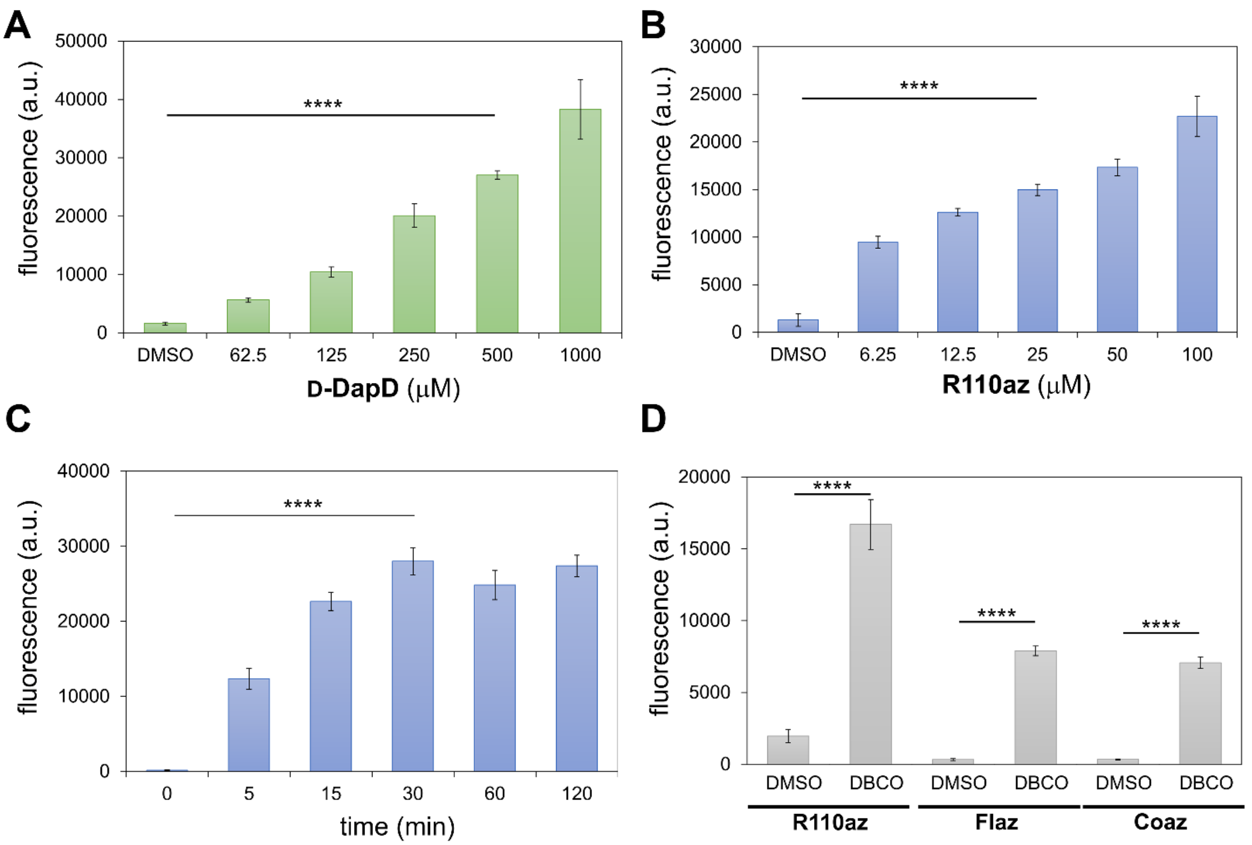


**Figure S3. *in vitro* PAC-MAN optimization.** A) Flow cytometry analysis of *S. aureus* incubated over-night with increasing concentrations of D-DapD, followed by incubation with 25 μM R110az for 30 min. B) Flow cytometry analysis of *S. aureus* incubated over-night with 500 μM D-DapD, followed by incubation with increasing concentrations of R110az for 30 min. C) Flow cytometry analysis of *S. aureus* incubated over-night with 500 μM D-DapD, followed by incubation with 25 μM R110az over time. D) Flow cyometry analysis of *S. aureus* incubated overnight with 500 μM D-DapD, followed by incubation with R110az, 6-azido-fluorescein (Flaz), and 3-azido-7-hydroxy coumarin (Coaz). 10000 events were recorded for each condition. Data are represented as mean ± SD of biological replicates (n= 3). *P*-values were determined by a two-tailed t-test (***$p < 0.001$, ****$p < 0.0001$, ns = not significant).

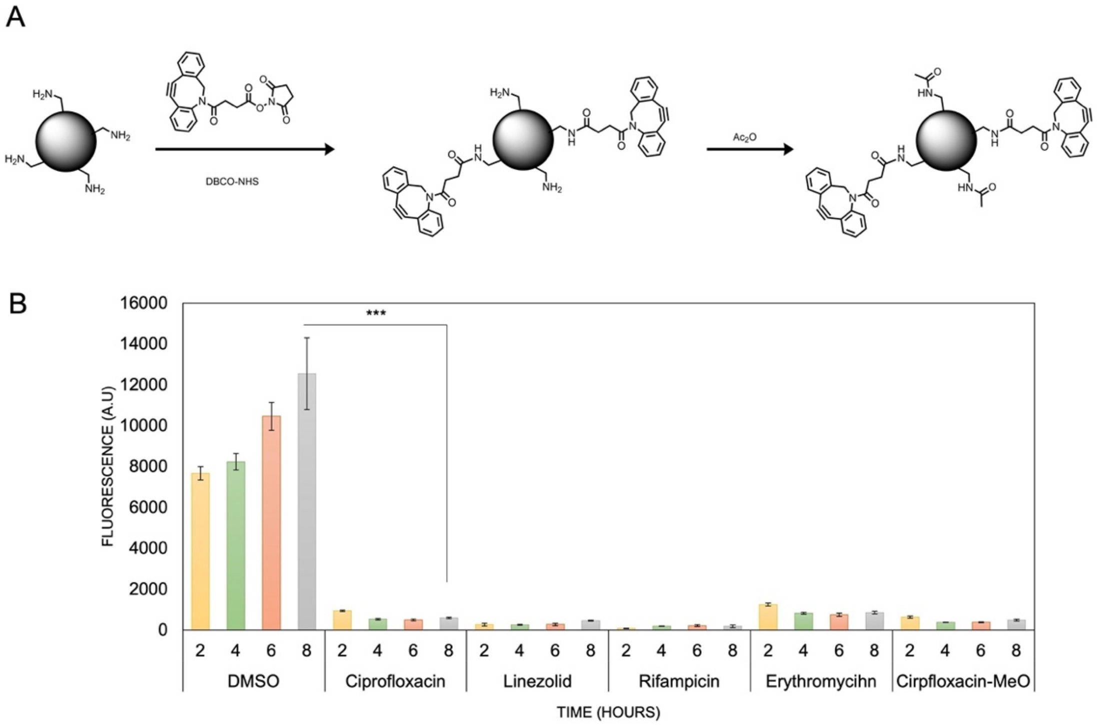


**Figure S4. Synthesis of DBCO- modified polystyrene beads and azAbx competition to DBCO- modified polystyrene beads.** A) Amine functionalized polystyrene beads were reacted with DBCO-NHS to form DBCO functionalized flow cytometer beads. The beads were further reacted with acetic anhydride to cap the remaining free amines. B) DBCO functionalized beads were incubated with 25 μM of azAbx over time, followed by incubation with 25 μM of Flaz for 30 min. Data are represented as mean ± SD of biological replicates (n= 3). 10000 events were recorded for each condition. *P*-values were determined by a two-tailed t-test (***p < 0.001, ****p < 0.0001, ns = not significant).

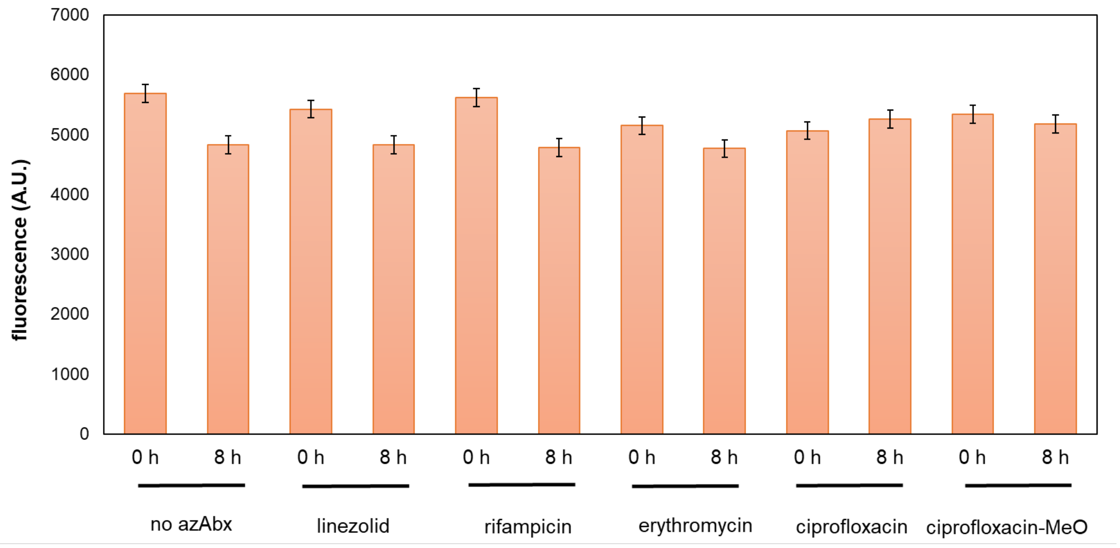


**Figure S5. Non-azide Abx competition to DBCO-modified *S. aureus*.** *S. aureus* cells were incubated with 500 μM of D-DapD overnight followed by incubation with 25 μM of unmodified (non-azide) Abx for indicated time points. Following the unmodified Abx incubation, the cells were resuspended in 25 μM of R110az for 30 min and analyzed by flow cytometry. 10000 events were recorded for each condition. Data are represented as mean ± SD of biological replicates (n= 3).

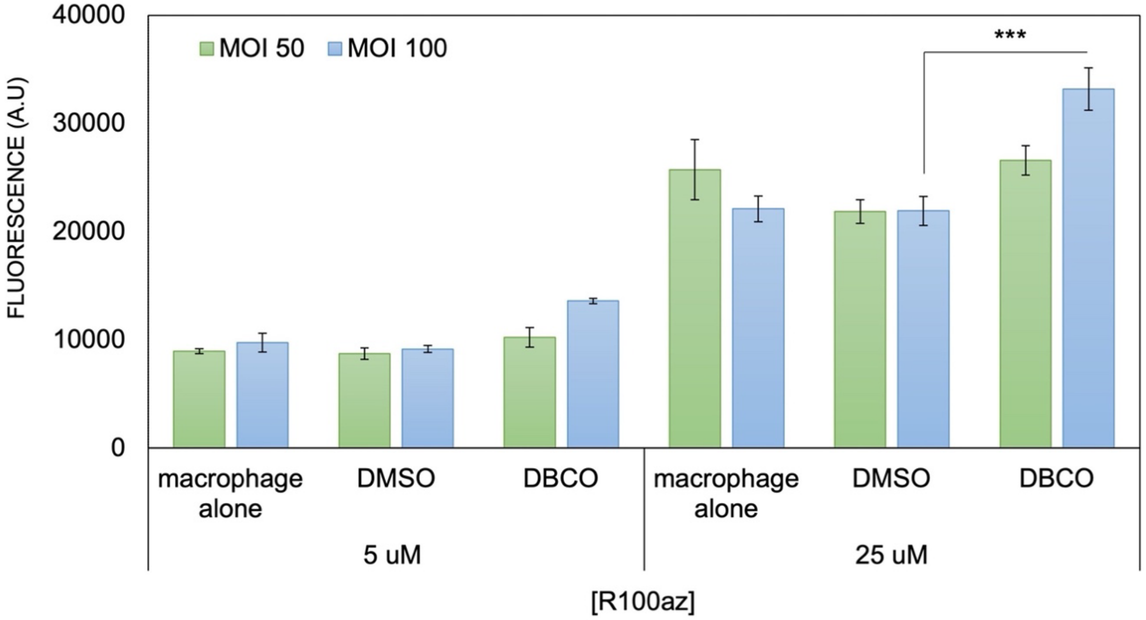


**Figure S6. Multiplicity of Infection (MOI) Assay.** *S. aureus* was grown overnight with 500 μM of D-DapD and infected into J774A macrophages at MOI 50 and MOI 100. Following bacterial uptake, varying concentrations of R110az were incubated for 30 min, and cells were analyzed by flow cytometry. 10000 events were recorded for each condition. Data are represented as mean ± SD of biological replicates (n= 3). *P*-values were determined by a two-tailed t-test (***p < 0.001, ****p < 0.0001, ns = not significant).

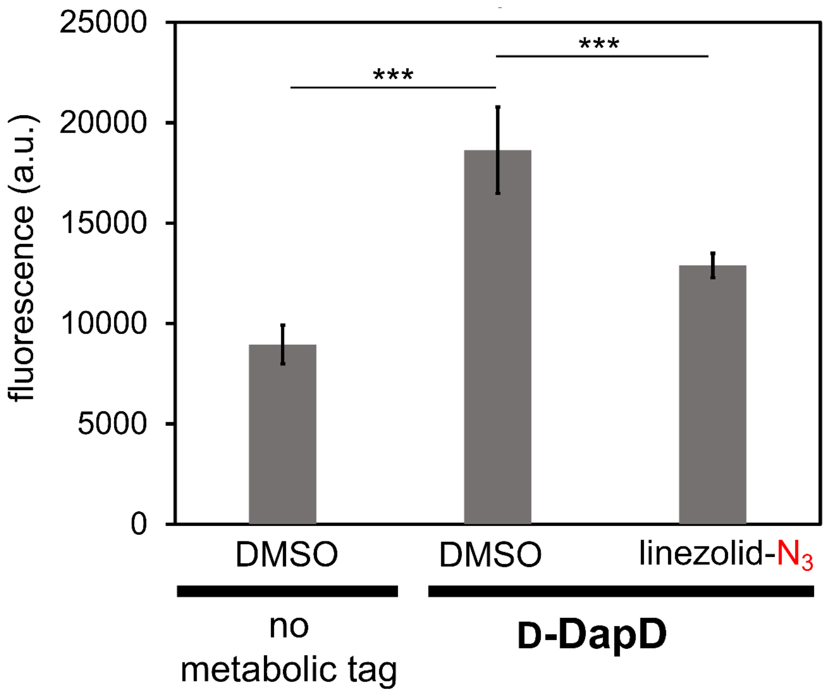


**Figure S7. Application of Assay Workflow to *S. pyogenes*.** A) Flow cytometry analysis of *S. pyogenes* incubated over-night with D-DapD or DMSO, followed by incubation with either DMSO or linezolid-$N_3$, and then incubated with 25 μM R110az for 30 min. B) *S. pyogenes* was grown overnight with 500 μM of D-DapD and infected into J774A macrophages at MOI 100. Following bacterial uptake, DMSO or linezolid-$N_3$ was incubated with the macrophages for 2 hours. The cells were then treated with R110az for 30 min, and cells were analyzed by flow cytometry. 10000 events were recorded for each condition. Data are represented as mean ± SD of biological replicates (n= 3). *P*-values were determined by a two-tailed t-test (***p < 0.001, ****p < 0.0001, ns = not significant).

| COMPOUND | MIC (mg/l) | COMPOUND | MIC (mg/l) |
|---|---|---|---|
| **CIPROFLOXACIN-$N_3$** | 16 | **CIPROFLOXACIN** | 2 [a] |
| **LINEZOLID-$N_3$** | 16 | **LINEZOLID** | 4 [b] |
| **RIFAMPICIN-$N_3$** | 2 | **RIFAMPICIN** | 0.1 [c] |
| **ERYTHROMYCIN-$N_3$** | >32 | **ERYTHROMYCIN** | 0.4 [c] |

**Table S1. Minimum Inhibitory Concentration (MIC) of azAbx library.** *S. aureus* 25923 was grown overnight and diluted with TSB to $OD_{600}$ = 0.2 and was regrown to $OD_{600}$ = 1.0. Cultures were diluted to $10^6$ CFU/mL in cation-adjusted MH broth. 100 $\mu$L was inoculated into each well of a U-bottom 96-well plate containing 100 $\mu$L of serially diluted azABx solution. After incubation for 24 hours at 37$^o$C, MIC values were determined by the lowest concentration of azAbx resulting in no bacterial growth visible to the naked eye based on three biological replicates.

[a] Antibiotics (Basel). 2021 Sep 24;10(10):1159. doi: 10.3390/antibiotics10101159.

[b] Antimicrob Agents Chemother. 1996 Apr;40(4):839-45. doi: 10.1128/AAC.40.4.839.

[c] Antimicrob Agents Chemother. 1981 Jun;19(6):1050-5. doi: 10.1128/AAC.19.6.1050.

**Materials.**

All peptide related reagents and protected amino acids were purchased from Chem-Impex. DBCO-NHS (Catelog # BP-22231) was purchased from Broad Pharm. Deacetamide Linezolid Azide was purchased from Toronto Research Chemicals (Cat #: D195600). Dulbecco's Modified Eagle's Medium (DMEM) was purchased from VWR. Fetal Bovine Serum (FBS) was purchased from R&D Systems. Penicillin-Streptomycin was purchased from Sigma-Aldrich. All other organic chemical reagents were purchased from Fisher Scientific or Sigma Aldrich and used without further purification.

**Methods.**

**Bacterial Cell Culture.** Bacterial cells were cultured in specified media in an aerobic environment while shaking at 250 rpm at 37$^{o}$C. *Staphylococcus aureus* (*S. aureus*) ATCC 25923 and *S. aureus* USA 300 were grown in Tryptic Soy Broth (TSB). *Streptococcus pyogenes* (*S. pyogenes*) ATCC 49399 were grown in Brain Heart Infusion (BHI) media. BLS2 organisms should be manipulated using proper protective equipment.

**Mammalian Cell Culture.** J774A.1 cells were cultured in Dulbecco's modified Eagle's medium (DMEM) supplemented with 10% (v/v) FBS, 50 IU/mL penicillin, 50 ug/mL streptomycin, and 2 mM L-glutamine in a humidified atmosphere of 55 $CO_2$ at 37$^{o}$C.

**Labeling of Whole Bacterial Cells with D-DapD.** *S. aureus* was grown over-night to stationary phase in TSB while shaking (250 rpm) at 37$^{o}$C. Bacterial cells from the overnight growth were used to inoculate TSB (1:100) supplemented with 500 μM D-DapD (or indicated concentration) and incubated at 37$^{o}$C with shaking (250 rpm) for 16 h. The bacteria were harvested, washed three times with 1X phosphate buffered saline (PBS). The bacterial cells were resuspended in 1X PBS supplemented with 25 μM (or indicated concentration) of either 6-azido-rhodamine 110 (R110az, Lumiprobe #D5230), 6-azido-fluorescein (Flaz, Lumiprobe #D1530), or 3-azido-7-hydroxy coumarin (Coaz, Sigma Aldrich #909513) in 1X PBS and incubated at 37$^{o}$C for 30 min. The bacteria were spun down to remove excess dye, immediately fixed with a 2% formaldehyde solution in 1X PBS, and analyzed using the Attune NxT Flow Cytometer (Thermo Fisher) equipped with a 488 nm laser with 530/30 nm bandpass filter. The data were analyzed using Attune Nxt software. The same procedure was followed for *S. pyogenes* grown in BHI.

**Sacculi Isolation of D-DapD labeled *S. aureus*.** *S. aureus* ATCC 25923 was grown over-night to stationary phase in TSB medium. A 2 mL culture volume containing either DMSO or 500 μM **D-DapD** in TSB medium was inoculated (1:100) from the stationary phase cultures and allowed to grow for 16 h while shaking (250 rpm) 37$^{o}$C. The cultures were harvested, resuspended in 25 μM of R110az in 1X PBS for 30 min at 37$^{o}$C, and washed thrice with 1X PBS. The samples were boiled at 100 $^{o}$C for 25 min and centrifuged at 14,000 g for 5 min at 4$^{o}$C. The cells were placed in 2 mL of 2% (w/v) sodium dodecyl sulfate (SDS) and boiled for 30 min followed by centrifugation at 14,000 g for 5 min at 4$^{o}$C. Cells were then washed 6 times with DI water to remove the SDS. After washing, cells were resuspended in 2 mL of 20 mM Tris buffer (pH 8.0). Pellets were treated with

800 μg DNase for 24 h followed by 800 μg trypsin for another 24 h at 37°C while shaking (115 rpm). Pellets were boiled for 25 min followed by centrifugation at 14,000 g for 5 min at 4°C. The pellet was harvested by centrifugation at 16,000 g for 5 min, resuspended in 1X PBS, and further diluted for analysis by flow cytometry and confocal imaging.

**Azide Antibiotic (azAbx) Competition of D-DapD Labeled Whole Bacterial Cells.** *S. aureus* was grown over-night to stationary phase in TSB while shaking (250 rpm) at 37°C. Bacterial cells from the overnight growth were used to inoculate TSB (1:100) supplemented with 500 µM **D-DapD** and incubated at 37°C with shaking (250 rpm) for 16 h. The bacteria were harvested, washed three times with 1X PBS. The bacterial cells were resuspended in 1X PBS supplemented with 25 μM of azide- Abx for the indicated amount of time at 37°C. The bacteria were spun down to remove excess antibiotics and were resuspended in 1X PBS supplemented with 25 μM 6-azido-rhodamine 110 (R110az, Lumiprobe #D5230) in 1X PBS and incubated at 37°C for 30 min. The bacteria were spun down to remove excess dye, immediately fixed with a 2% formaldehyde solution in 1X PBS, and analyzed using the Attune NxT Flow Cytometer (Thermo Fisher) equipped with a 488 nm laser with 530/30 nm bandpass filter. The data were analyzed using Attune Nxt software.

**Non-Azide Antibiotic Competition of D-DapD Labeled Whole Bacterial Cells.** *S. aureus* was grown over-night to stationary phase in TSB while shaking (250 rpm) at 37°C. Bacterial cells from the overnight growth were used to inoculate TSB (1:100) supplemented with 500 µM **D-DapD** and incubated at 37°C with shaking (250 rpm) for 16 h. The bacteria were harvested, washed three times with 1X PBS. The bacterial cells were resuspended in 1X PBS supplemented with 25 μM of non-azide abx for the indicated amount of time at 37°C. The bacteria were spun down to remove excess antibiotics and were resuspended in 1X PBS supplemented with 25 μM 6-azido-rhodamine 110 (R110az, Lumiprobe #D5230) in 1X PBS and incubated at 37°C for 30 min. The bacteria were spun down to remove excess dye, immediately fixed with a 2% formaldehyde solution in 1X PBS, and analyzed using the Attune NxT Flow Cytometer (Thermo Fisher) equipped with a 488 nm laser with 530/30 nm bandpass filter. The data were analyzed using Attune Nxt software.

**Minimum Inhibitory Concentration (MIC) Assay for azAbx against *S. aureus.*** Compounds were serially diluted 2-fold to yield 11 test concentrations. *S. aureus* was grown in an overnight culture and diluted the next day to optical density 600 ($OD_{600}$) of 0.2 and regrown to $OD_{600}$ of 1. All cultures were diluted to $10^6$ CFU/mL in cation-adjusted Muller Hilton broth and 100 μL was inoculated into each well of a U-bottom 96-well plate containing 100 μL of compound solution. Plates were grown statically at 37°C for 24 hours upon which time wells were visually evaluated for bacterial growth. The MIC was determined by the lowest concentration of compounds resulting in no bacterial growth visible to the naked eye, based on the majority of three independent experiments.

**Confocal Microscopy Analysis of Whole Bacterial Cells and Bacterial Sacculi.** Glass microscope slides were spotted with a 1% agarose pad and 2 μL of the fixed bacterial samples were deposited onto the agarose. Samples were covered with a micro cover glass and imaged using a Zeiss 880/990 multiphoton Airyscan microscopy system (63x oil-immersion lens) equipped with a 488 nm laser. Images were obtained and analyzed via Zeiss Zen software. We acknowledge the Keck Center for Cellular Imaging and for the usage of the Zeiss 880/980 multiphoton Airyscan microscopy system (PI- AP: NIH-OD025156).

**DBCO Modification of Polystyrene Beads.** 100 μL amino functionalized polystyrene beads (5% w/v, 5 mg) were spun down at 21000 g for 10 min in a 1.7 mL ebb tube and washed with 1 mL deionized water before use. The beads were then spun down and resuspended in 1 mL 20 mM sodium borate buffer pH 9 with 2 μg/mL DBCO-NHS and reacted in 37°Cfor 2 h with shaking. The resulting beads were spun down at 21,000 g for 10 min, washed once and resuspended in sodium borate buffer. 20 μL acetic anhydride was added to the suspension and reacted in 37°C for 2h with shaking. The resulting product was then spun down and washed twice with 1 mL PBS and resuspend in 1 mL PBS for further use. Control beads were acetylated with acetic anhydride directly after wash with deionized water in the same conditions.

**Competition with azAbx to DBCO Modified Polystyrene Beads.** DBCO modified and acetylated polystyrene beads were 1 to 1 diluted in PBS before adding to the assay. To a 96-well plate added 5 μL beads each well to 25 μM of each azide-abx in the library with a final volume of 100 μL in multiplicity of 12. The plate was incubated in 37°C for 2 h, 4 h, 6 h, 8 h. At each time point, 3 wells of the beads in each group were transferred to a 0.45 mm MultiScreen HV Filter Plate (Sigma, Cat # MSHVN4510) and vacuum filtered. The beads were then washed with 200 mL PBS two times to remove the residue azide-abx and resuspended in 100 μL PBS until next reaction. After collection of the last time point, 100 μL of 25 mM **Fl-Az** was added to each well and the plate was then incubated in 37℃ for 30 min. The samples were then vacuum filtered and washed twice with 200 μL PBS and then resuspended in 200 μL PBS. The samples were then analyzed by Attune™ NxT Flow Cytometer (Thermo Fisher) equipped with a 488 nm laser with 530/30 nm bandpass filter. The data were analyzed using Attune Nxt software.

**azAbx Competition to D-DapD Labeled Intracellular *S. aureus*.** *S. aureus* ATCC 25923 was grown to stationary phase in TSB while shaking (250 rpm) at 37 °C. Bacterial cells from the overnight growth were used to inoculate TSB (1:100) supplemented with 500 μM **D-DapD** and incubated at 37 °C with shaking (250 rpm) for 16 h. The bacteria were harvested, washed three times with 1X PBS. The bacterial cells were resuspended with 1X PBS to the original culture volume. J774A.1 cells were cultured as described above. On the day prior to the experiment, J774A.1 cells were seeded into a 48- well plate and allowed to adhere. On the day of the experiment, J774A.1 cells were washed with 1X PBS by centrifuging 5 min at 100 *g*. The washed J774A cells were then mixed with **D-DapD** labeled *S. aureus* (MOI 100) in DMEM + 10% FBS containing no antibiotics. The cell mixture was then incubated at 37 °C for 1 hour to induce phagocytosis. The cell mixture was centrifuged for 5 min at 100 *g* and media was replaced with DMEM + 10% FBS + 300 μg/mL gentamycin and incubated at 4°C for 30 min. The cells were washed three times with 1X PBS and incubated with either a solution containing 25 μM R110az (for no Abx competition samples) or 25 μM azide- Abx + 300 μg/mL gentamycin in 1X

PBS for indicated amounts of time at 37°C. The Abx medium was removed, and the cells were incubated with a solution of 25 μM R110az in 1X PBS for 30 min at 37°C.The cells were washed thrice with 1X PBS and fixed for 30 min with 4% formaldehyde in 1X PBS. Samples were then removed from the well plate by scraping and analyzed *via* flow cytometry as described above. The same procedure was followed for *S. pyogenes* grown in BHI.

**Confocal Analysis of D-DapD Labeled Intracellular *S. aureus*.** *S. aureus* was grown over-night to stationary phase in TSB while shaking (250 rpm) at 37 °C. Bacterial cells from the overnight growth were used to inoculate TSB (1:100) supplemented with 500 μM **D-DapD** and incubated at 37 °C with shaking (250 rpm) for 16 h. The bacteria were harvested, washed three times with 1X PBS. The bacterial cells were resuspended in 1X PBS supplemented with 25 μM of R110az in 1X PBS and incubated at 37°C for 30 min. The bacteria were washed thrice and resuspended in in 1X PBS to the original culture volume. J774A.1 cells were cultured as described above. On the day prior to the experiment, J774A.1 cells were seeded into 35 mm glass bottom microwell dishes and allowed to adhere. On the day of the experiment, J774A.1 cells were washed with 1X PBS by centrifuging 5 min at 100 *g*. The washed J774A cells were then mixed with **D-DapD** labeled *S. aureus* (MOI 100) in DMEM + 10% FBS containing no antibiotics. The cell mixture was then incubated at 37°C for 1 hour to induce phagocytosis. The cell mixture was centrifuged for 5 min at 100 *g* and media was replaced with DMEM + 10% FBS + 300 μg/mL gentamycin and incubated at 4°C for 30 min. The cells were washed thrice with 1X PBS and fixed for 30 min with 4% formaldehyde in 1X PBS. J774A.1 macrophages were then treated with 5 μg/mL of TMR-tagged Wheat Germ Agglutinin (Vector Laboratories, RL-1022) for 30 min at 4°C and imaged using a Zeiss 880/990 multiphoton Airyscan microscopy system (63x oil-immersion lens) equipped with 488 nm and 550 nm lasers. Images were obtained and analyzed via Zeiss Zen software. We acknowledge the Keck Center for Cellular Imaging and for the usage of the Zeiss 880/980 multiphoton Airyscan microscopy system (PI- AP: NIH-OD025156).

**Synthesis and Characterization.**

**Scheme S1.** Synthesis of D-DapD

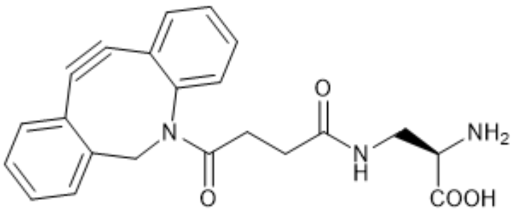


To a 25 mL peptide synthesis vessel with 100 mg 2-Chlorotrityl chloride resin (0.142 mmol) resuspended in 15 mL dry dichloromethane, was added Nα-Boc-Nβ-Fmoc-D-2,3-diaminopropionic acid (D-Dap, 67 mg, 1.1 eq, 0.16 mmol), and DIEA (4.4 eq, 0.11 mL, 0.62 mmol). The resin was shaken for 1 hour at room temperature and washed with methanol and dichloromethane (3 times and 15 mL each). Fmoc protecting group was removed with 6M piperazine in N, N-Dimethylformamide (DMF, 15 mL) for 30 min at room temperature and washed as before. DBCO was coupled on the side chain of D-Dap on resin. 25-30 mg DBCO-NHS was dissolved in 1 mL dry DMF and added to the 25 mL peptide synthesis vessel with 100 mg equivalent 2-Chlorotrityl chloride resin with D-Dap resuspended in 2 mL DMF. The resin was shaken overnight at room temperature and washed with methanol and dichloromethane (3 times and 15 mL each). The resin was then added 20% trifluoroacetic acid (TFA) in dichloromethane after wash and shaken in room temperature for 1 h. The liquid phase was filtered and concentrated with nitrogen flow and added icy ether to precipitate the peptide. The ether layer was decanted, and the resulting solid was washed with icy ether and air dried. The crude material was purified with reverse phased high performance liquid chromatography (RP-HPLC) using a 40 to 100% linear gradient of methanol in $H_2O$/MeOH with 0.1% TFA to yield **D-DapD**. The sample was analyzed for purity using a Waters 1525 with a Phenomenex Luna 5μ C8(2) 100 Å (250 x 4.6 mm) column; gradient elution with $H_2O/CH_3CN$.

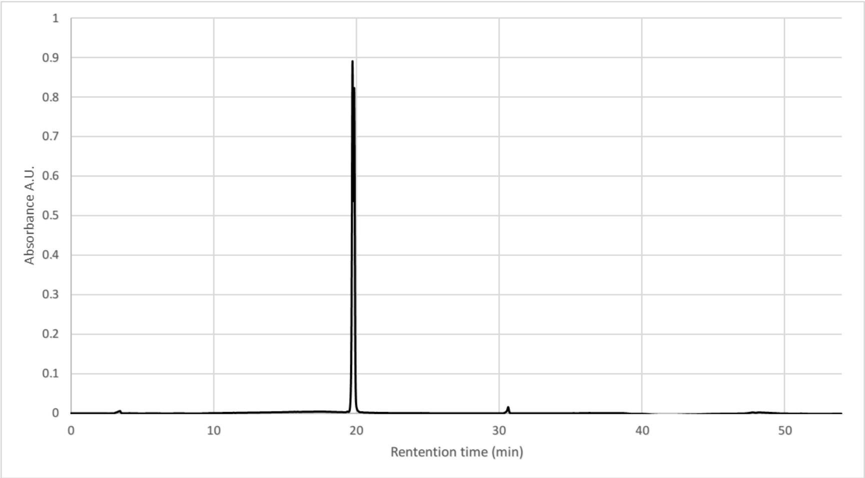


Molecular weight was confirmed using an Agilent LC-QTOF (Agilent 1260 Infinity II Prime LC with Agilent 6545B QTOF).

Calculated $(M+H)^+$, 392.1605, found: $(M+H)^+$ 392.1607.

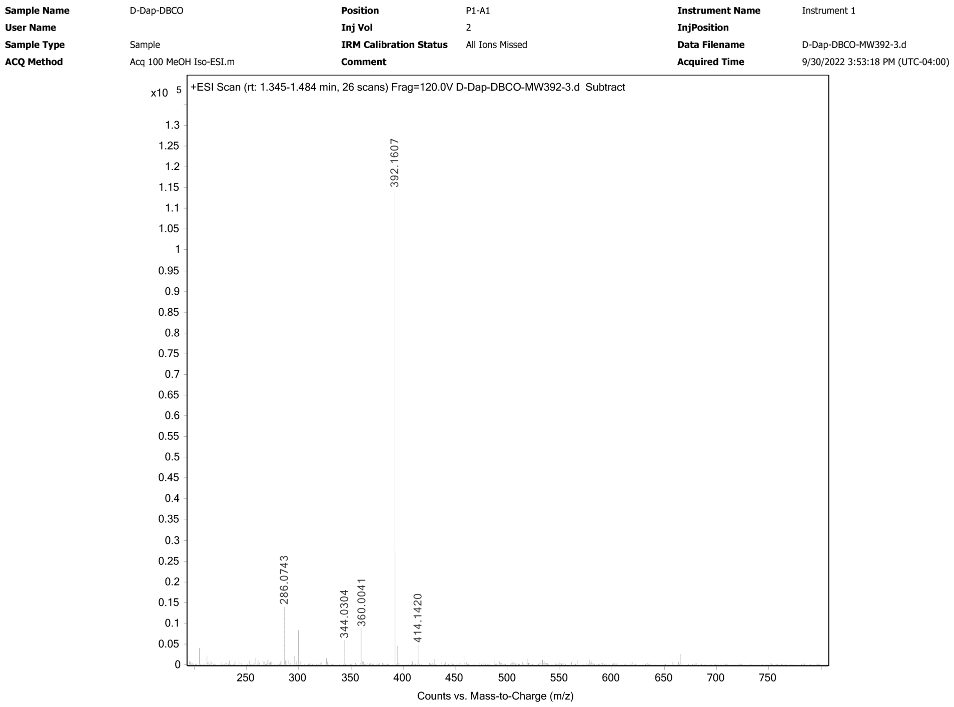


*Note: $^1$H and $^{13}$C-NMR spectra for all new compounds and intermediates for characterization were acquired on a Varian 600MHz spectrophotometer. All NMR spectra were analyzed using MestreNova software. Residual solvent signal from $CDCl_3$, $CD_3OD$ and DMSO-d6 referenced to tetramethylsilane (TMS) were used as reference standards for defining chemical shifts of $^{1}H$ or $^{13}C$ spectra of compounds. Chemical shifts are reported in δ ppm and coupling constants (J) are reported in Hertz [Hz]. Deuterated solvents were used as received from Cambridge Isotopes.

$^{1}H$-NMR

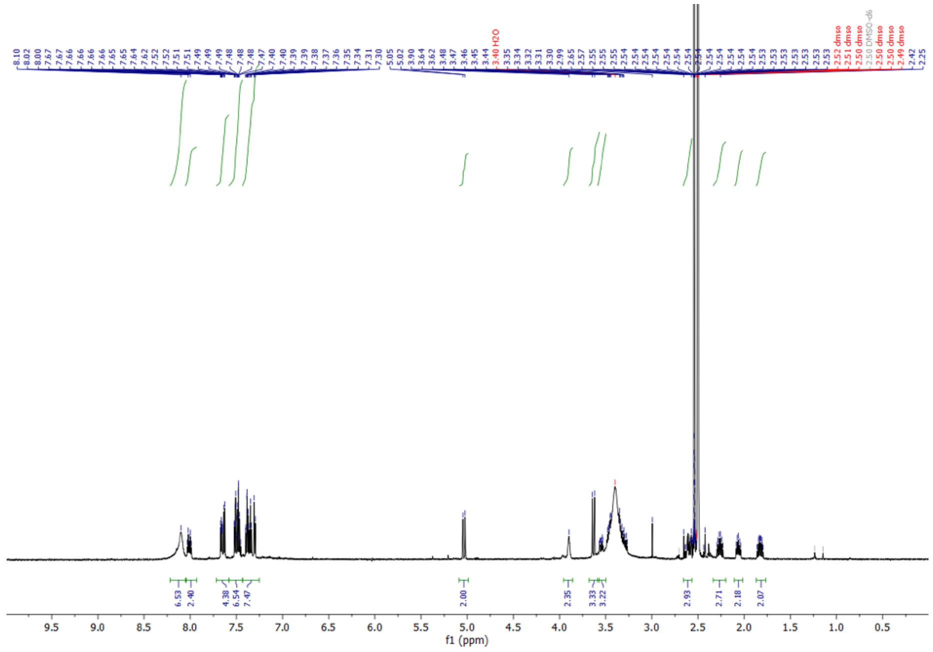


$^{13}C$-NMR

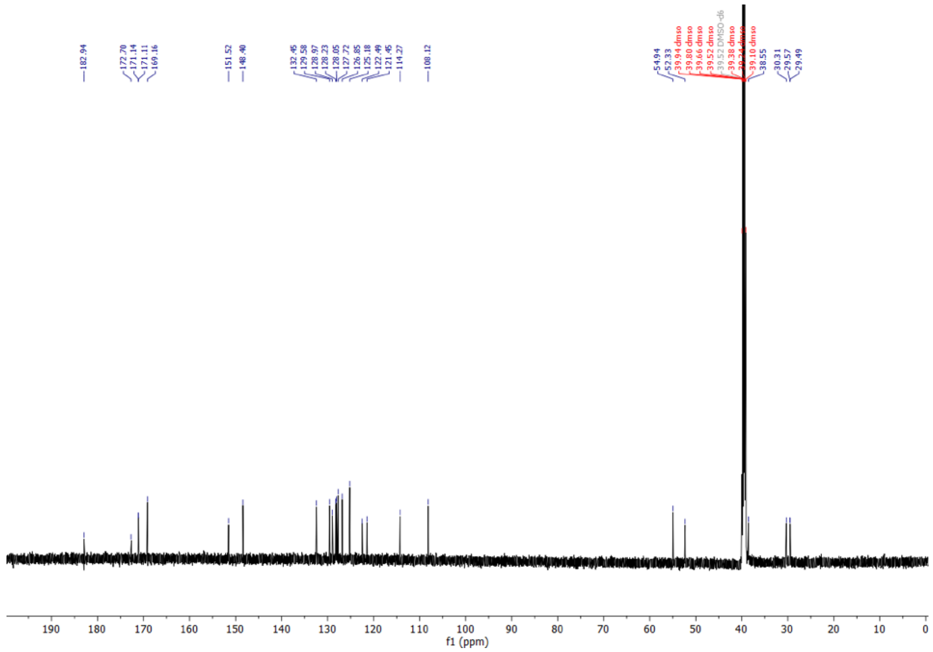

**Scheme S2.** Synthesis of Rifampicin-azidobutane

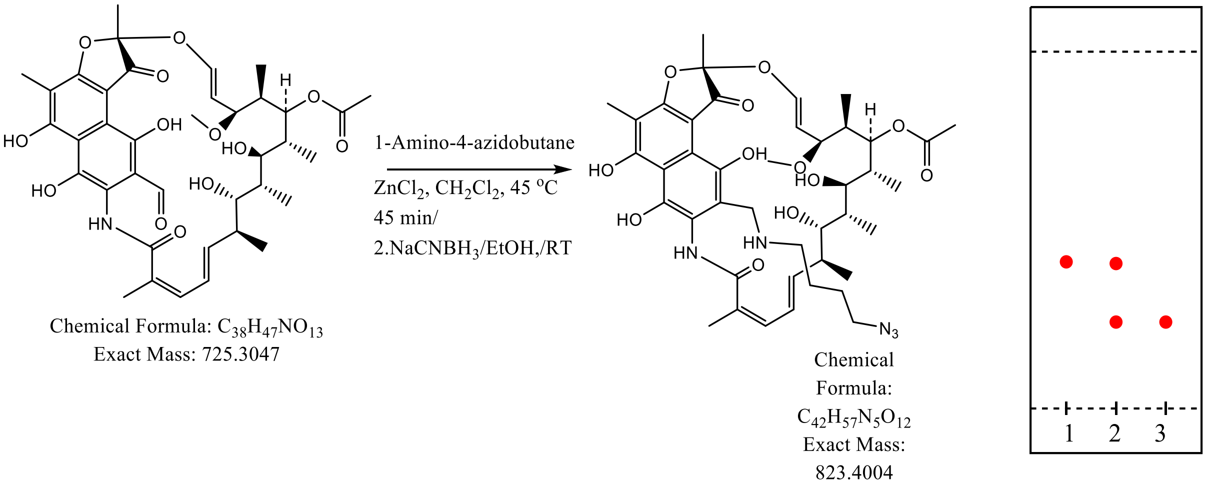


Rifamycine aldehyde (72.5mg. 0.1 mmol) dissolved in 5.0mL of anhydrous methylene chloride in 50 mL RB flask equipped with reflux condenser, to this solution was added 1-amino-4-azidobutane (17.1 mg, 0.15 mmol), followed by $ZnCl_2$ (11.0mg, at room temperature and the mixture was refluxed at 45 °C in an oil bath for 45 minutes. The mixture was the allowed cooled to room temperature. The solvents were evaporated, and crude material as is used for reduction of iminium. The crude material was dissolved in ethanol (200% proof, 2.5mL), sodium cyanoborohydride (10.7 mg, 0.17mmol) was added to it and stirred the mixture at room temperature for 2 hr. TLC analysis of crude material indicated complete conversion starting material (Rt ~0.4) to polar new compound (Rt ~0.2) in EtOAc:methanol (95:5) . The volatiles were removed under vacuo using rotary evaporator. The remaining residue was dissolved in EtOAc and purified by column chromatography over silica gel using gradient ranging from ethyl acetate to 3% methanol EtOAc. Fractions showing homogeneity on TLC were combined and concentrated under reduced pressure to yield red solid (65.1 mg, 79%). Further purification with reverse phase HPLC (80:20; Water;Methanol) afforded pure compound used for assay.

Molecular weight was confirmed using an Agilent LC-QTOF (Agilent 1260 Infinity II Prime LC with Agilent 6545B QTOF).

Calculated m/z,824.534, found m/z, 824.4131

| | | | | | |
|---|---|---|---|---|---|
| **Sample Name** | MDC Rifampicin azide | **Position** | P1-A4 | **Instrument Name** | Instrument 1 |
| **User Name** | | **Inj Vol** | 1 | **InjPosition** | |
| **Sample Type** | Sample | **IRM Calibration Status** | Success | **Data Filename** | MDC rifampicin azide 1.3ul injection.d |
| **ACQ Method** | Acq 80_20 MeOH_H2O Iso.m | **Comment** | | **Acquired Time** | 8/26/2022 2:37:15 PM (UTC-04:00) |

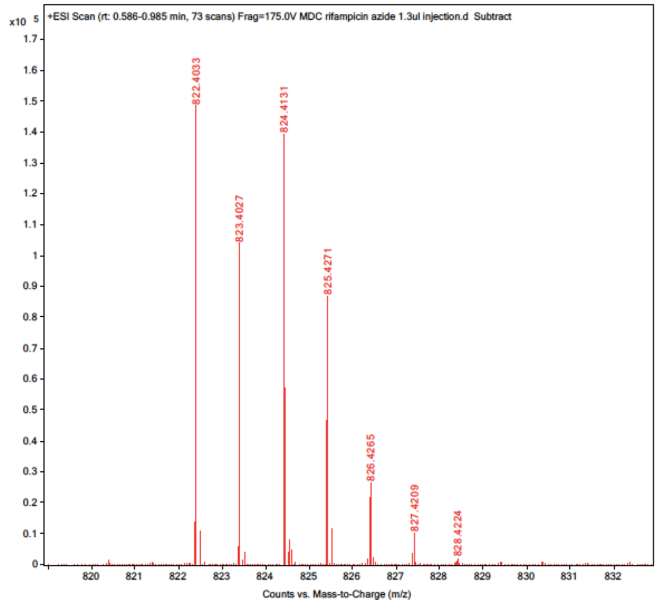


### $^{1}$H-NMR

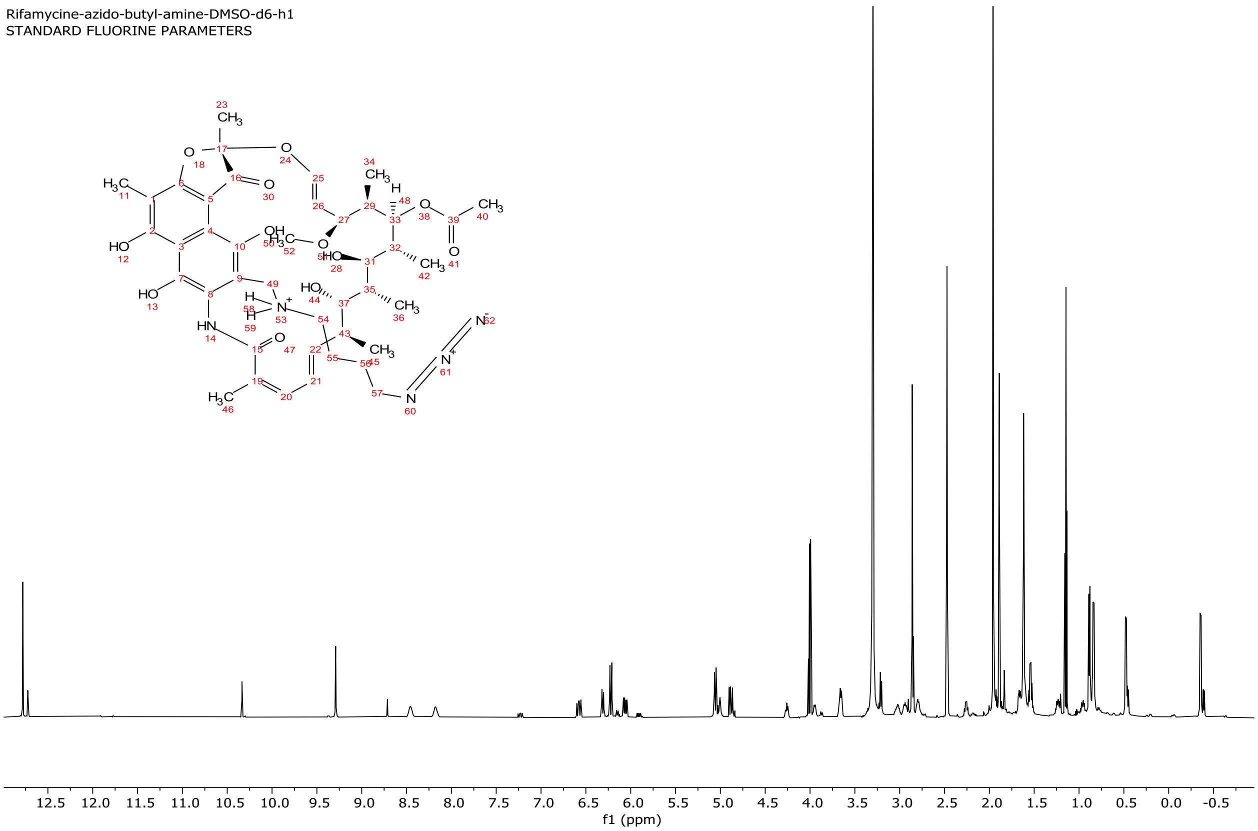

**Scheme S3.** Synthesis of Erythromycin-azidoacetamide

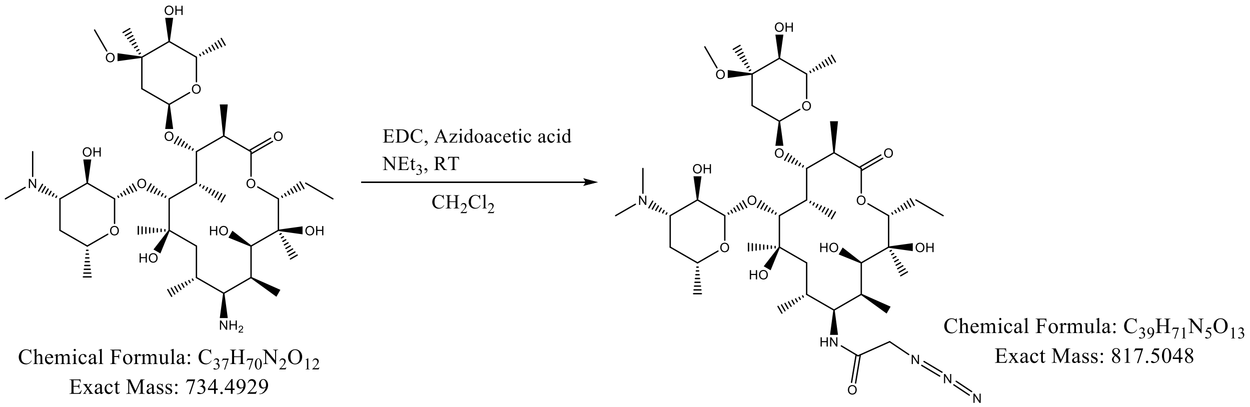


Erythromycin (25.1 mg, 0.034mmol) was dissolved in methylene chloride (5 mL), subsequently azido acetic acid (3.9 mg, 0.0386 mmol), EDC (11.5 mg) and triethyl amine (25.0ul) were added and mixture was stirred at room temperature for overnight (16hr). Next day, the solvents were evaporated under reduced pressure using rotary evaporator, the leftover crude material was purified by gradient reverse phase HPLC (C8, Water:MeOH; 95:5). The homogenous fractions collected at 205nm, were concentrated to yield pure compound.

Molecular weight was confirmed using an Agilent LC-QTOF (Agilent 1260 Infinity II Prime LC with Agilent 6545B QTOF).

Calculated $(M+D)^+$,819.5208, found $(M+D)^+$, 819.5321

| | | | | | |
|---|---|---|---|---|---|
| **Sample Name** | MDC erythromycin azide | **Position** | P1-A3 | **Instrument Name** | Instrument 1 |
| **User Name** | | **Inj Vol** | 1 | **InjPosition** | |
| **Sample Type** | Sample | **IRM Calibration Status** | Success | **Data Filename** | MDC erythro 1uL 8.26.22.d |
| **ACQ Method** | Acq 80_20 MeOH_H2O Iso.m | **Comment** | | **Acquired Time** | 8/26/2022 12:29:12 PM (UTC-04:00) |

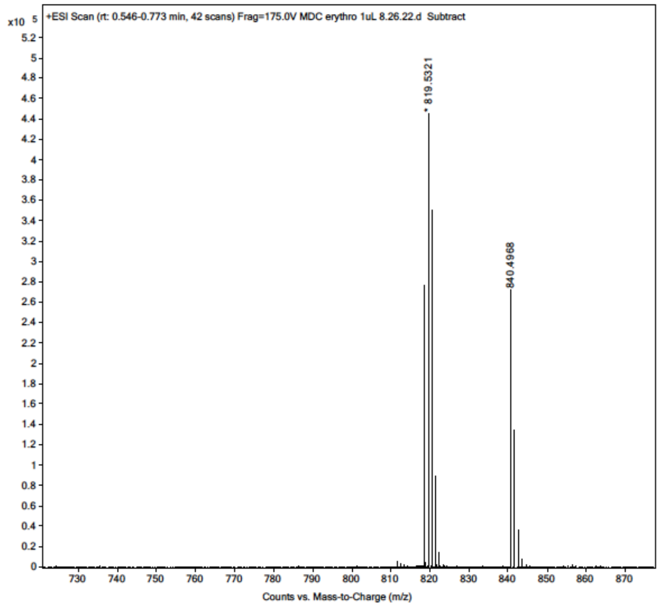


### [1]H-NMR

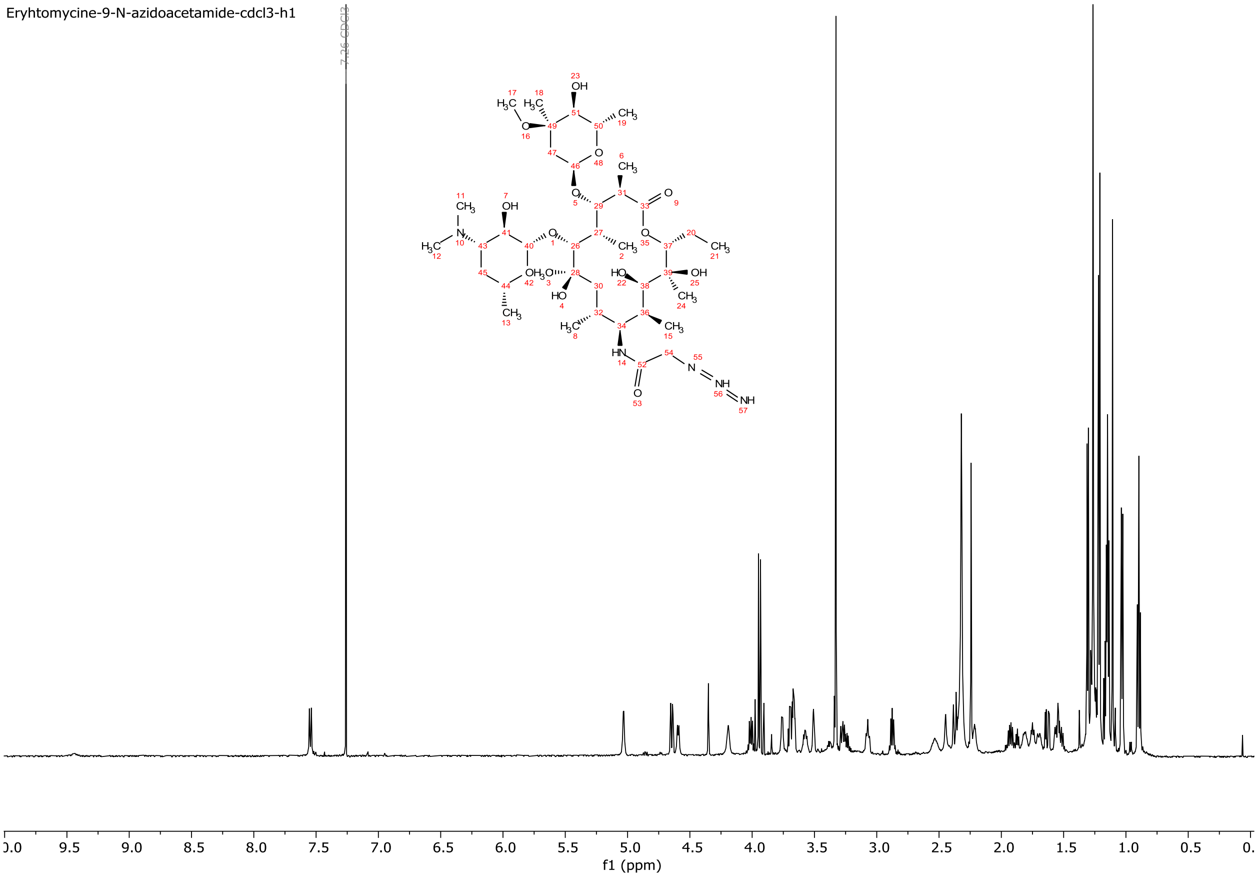

**Scheme S4.** Synthesis of N-azidoacetyl-Ciprofloxacin

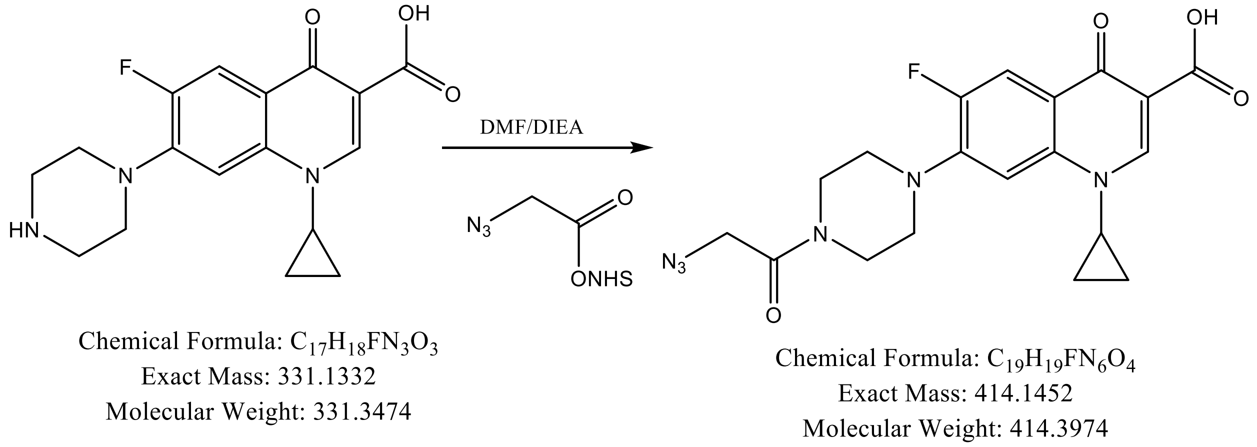


Ciprofloxacin (33.1mg, 0.1 mmol) suspended in anhydrous DMF (1.0mL), to which first DIEA (50uL) was added the stirred for minute prior to addition of Azido-N-hydroxy succinate (20.3mg, 0.102mmol) and the mixture was stirred at room temperature for overnight. Some white fine solid remained undissolved in the mixture. 15.0mL of ethyl acetate was added to reaction mixture followed by 10.0mL of water. The ethyl acetate layer was pale yellow colored while water layer was brownish colored. The ethyl acetate layer was separated, the aq. Layer was once more extracted with ethyl acetate 5.0mL. The combined ethyl acetate layer wash washed with brine, dried over $MgSO_4$ and concentrated on rotary evaporator under reduced pressure. The left-over residue was triturated with ether to obtained white solid. 1H-NMR of white solid indicated presence of unreacted azido acetic acid. The solid was dissolved in 0.5 mL DMSO and diluted with water to total volume of 10 mL. The analytical HPLC showed desired compound at Rt = min. Preparative HPLC using Water:$CH_3CN$ gradient (95:5 from 5 min. 25 min, 100% B for 5 min., total of 35 min run) purified to homogeneity by collected peak at Rt=23.4 min. (monitored at 280 and 320nm) as fractions. Combined fractions were concentrated first on rotary evaporator and then by lyophilized to obtain off white solid (14.7mg).

Molecular weight was confirmed using an Advion Expression® CMS mass spectrometer.

Calculated m/z, 414.4, found m/z, 415.2

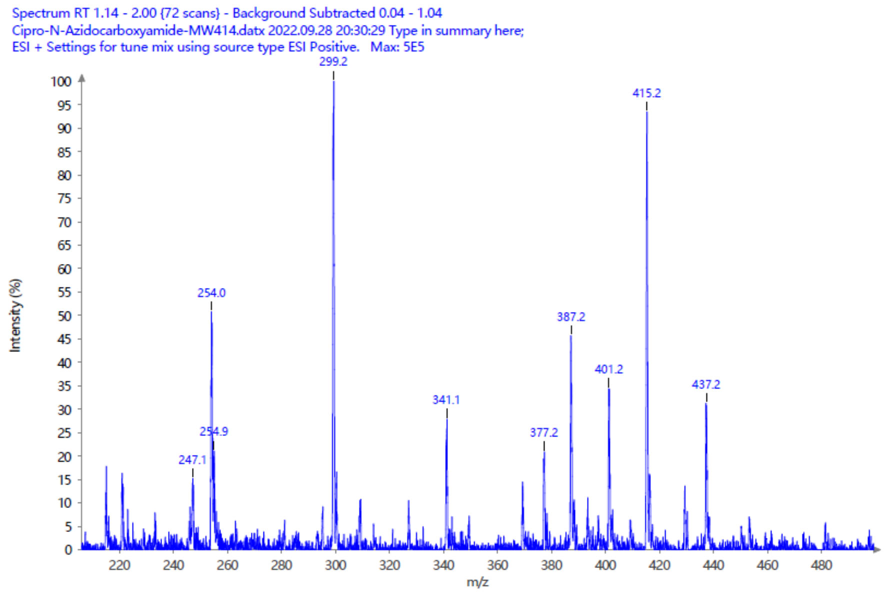


$^{1}$H-NMR

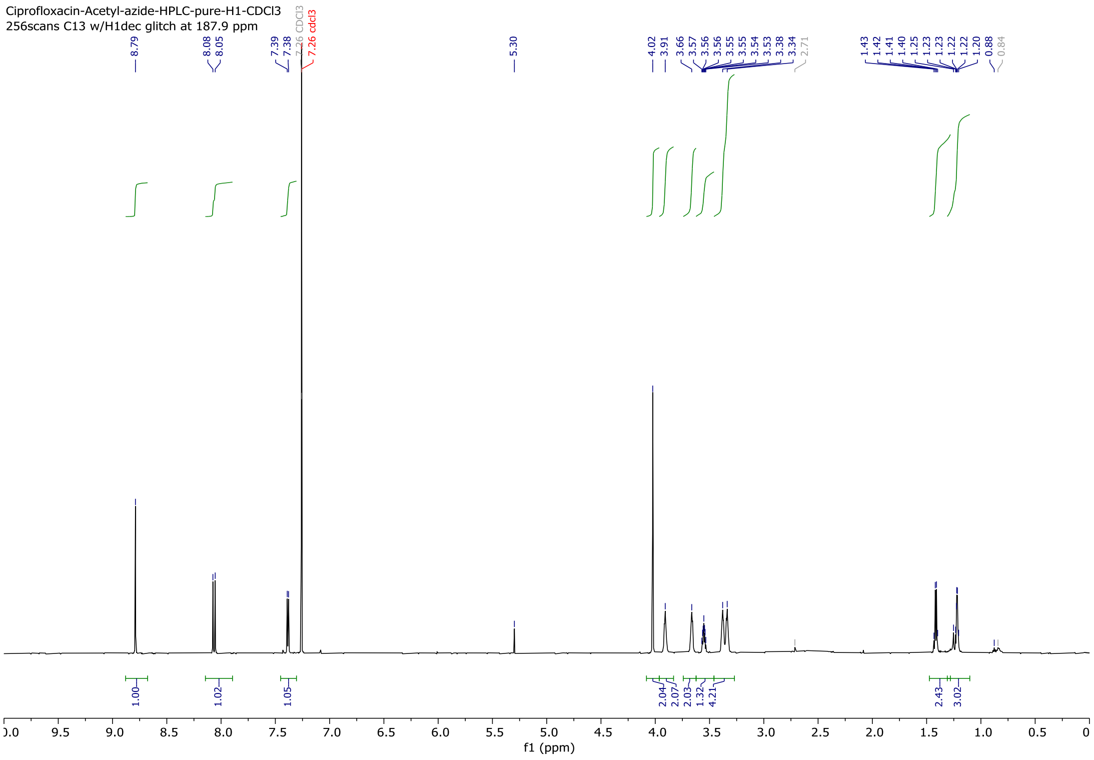

**Scheme S5.** Synthesis of Ciprofloxacin-azidoacetyl methyl ester

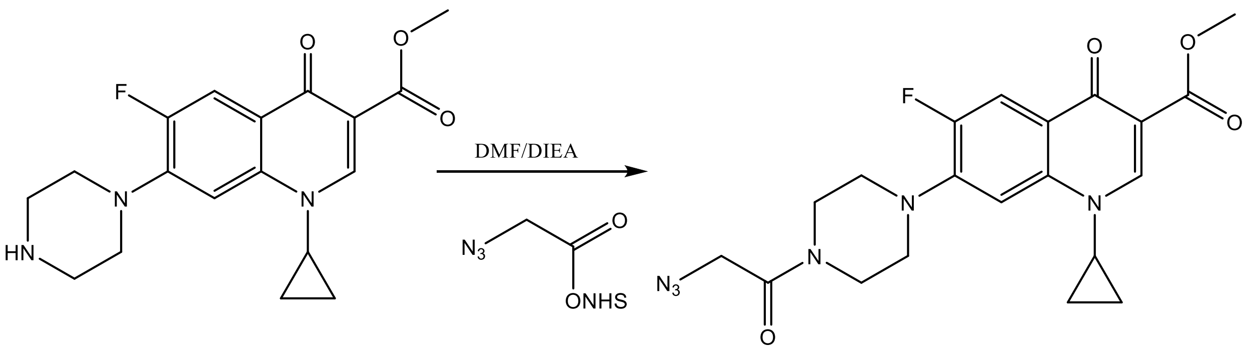


Chemical Formula: $C_{18}H_{20}FN_3O_3$
Exact Mass: 345.1489
Molecular Weight: 345.3744

Chemical Formula: $C_{20}H_{21}FN_6O_4$
Exact Mass: 428.1608
Molecular Weight: 428.4244

Ciprofloxacin-methyl ester as white solid was synthesized and characterized as per reported procedure using MeOH, $SOCl_2$. The ciprofloxacin-methyl ester (34.5 mg, 0.1mmol) was dissolved in anhydrous DMF (1.0 mL) and DIEA (50.0uL), followed by Azido-acetic acid NHS ester (25.3mg, 0.13 mmol) were added to it at room temperature and stirred for 4 hr. The TLC analysis indicated formation of new compound. The mixture was then diluted with chloroform (15.0mL), transferred to separatory funnel and washed thoroughly with water, and dil. HCl (0.1M). The organic layer was finally washed with brine, dried over $Na_2SO_4$ and concentrated under reduced pressure to yield crude material, which was purified with silica gel column chromatography using chloroform:methanol gradient (98:2), Homogenous fractions as analyzed by TLC were combined and concentrated on rotary evaporator, to afford white solid (32.5mg, 76%),

Molecular weight was confirmed using an Advion Expression® CMS mass spectrometer.

Calculated m/z, 428.4, found m/z, 429.2

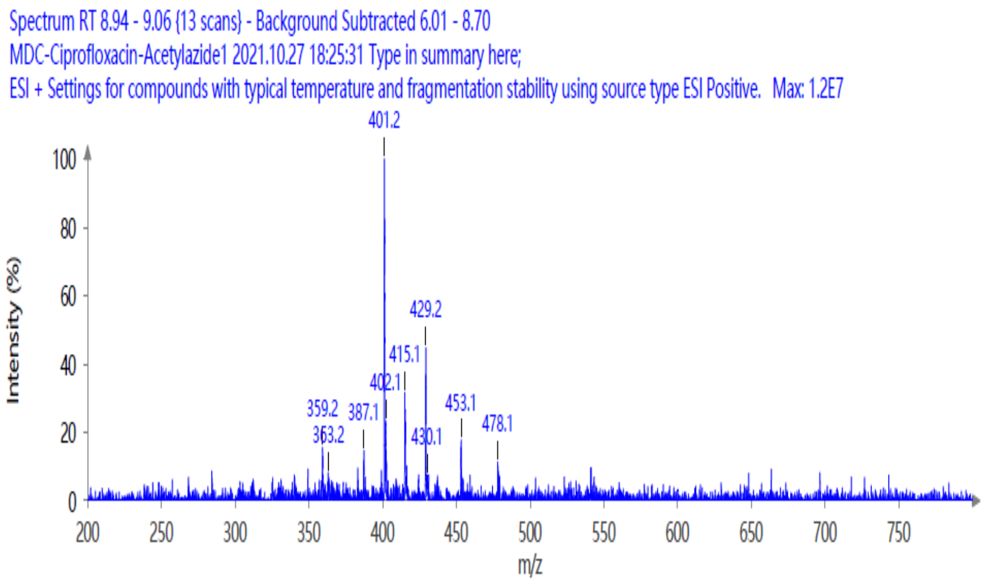


[1]H-NMR

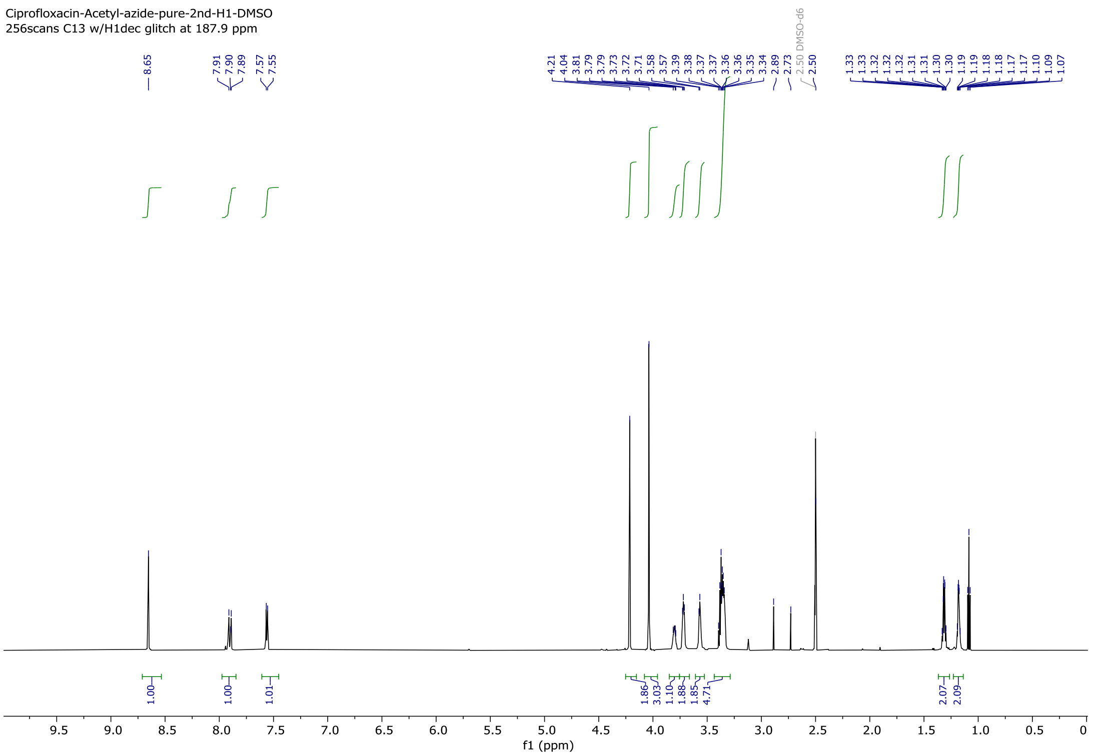
